# Structures of the Type IX Secretion/gliding motility motor from across the phylum *Bacteroidetes*

**DOI:** 10.1101/2022.01.28.478170

**Authors:** Rory Hennell James, Justin C. Deme, Alicia Hunter, Ben C. Berks, Susan M. Lea

## Abstract

Gliding motility using cell surface adhesins and export of proteins by the Type IX Secretion System (T9SS) are two phylum-specific features of the *Bacteroidetes*. Both of these processes are energized by the GldLM motor complex which transduces the protonmotive force at the inner membrane into mechanical work at the outer membrane. We previously used cryo-electron microscopy to solve the structure of the GldLM motor core from *Flavobacterium johnsoniae* at 3.9 Å resolution (*Nat Microbiol* (2021) **6:** 221-233). Here we present structures of homologous complexes from a range of pathogenic and environmental *Bacteroidetes* species at up to 3.0 Å resolution. These structures show that the architecture of the GldLM motor core is conserved across the *Bacteroidetes* phylum although there are species-specific differences at the N-terminus of GldL. The resolution improvements reveal a cage-like structure that ties together the membrane-proximal cytoplasmic region of GldL and influences gliding function. These findings add detail to our structural understanding of bacterial ion-driven motors that drive the T9SS and gliding motility.

**Importance:** Many bacteria in the *Bacteroidetes* phylum use the Type IX Secretion System to secrete proteins across their outer membrane. Most of these bacteria can also glide across surfaces using adhesin proteins that are propelled across the cell surface. Both secretion and gliding motility are driven by the GldLM protein complex which forms a nanoscale electrochemical motor. We used cryo-electron microscopy to study the structure of the GldLM protein complex from different species including the human pathogens *Porphyromonas gingivalis* and *Capnocytophaga canimorsus*. The organisation of the motor is conserved across species, but we find species-specific structural differences and resolve motor features at higher resolution. This work improves our understanding of the Type IX Secretion System, which is a virulence determinant in human and animal diseases.

## Introduction

The Type IX Secretion System (T9SS) is a protein export system found exclusively in the *Bacteroidetes* phylum of Gram-negative bacteria (1, 2). Substrates of the T9SS are transported to the periplasm by the Sec system following which a conserved C-terminal domain (CTD) directs export through an outer membrane T9SS translocon (Figure 1a) (2, 3). In most cases the CTD is then removed and the substrate protein is either released into the environment or anchored to the outer membrane as a lipoprotein (4). The human oral pathogen *Porphyromonas gingivalis* uses the T9SS to secrete gingipain proteases and other virulence factors to evade the host immune system (5). The T9SS has also been identified as essential to the virulence of several economically-relevant fish and poultry pathogens (6–8). In commensal and environmental *Bacteroidetes* the T9SS is characteristically used to secrete enzymes that enable the organisms to utilise complex polysaccharides as a food source (1, 9, 10). Many *Bacteroidetes* with a T9SS also exhibit gliding motility in which cells travel rapidly across surfaces (2). This motility depends on the movement of cell surface adhesin molecules which are secreted to the cell surface by the T9SS (Figure 1a).

**Figure 1.**
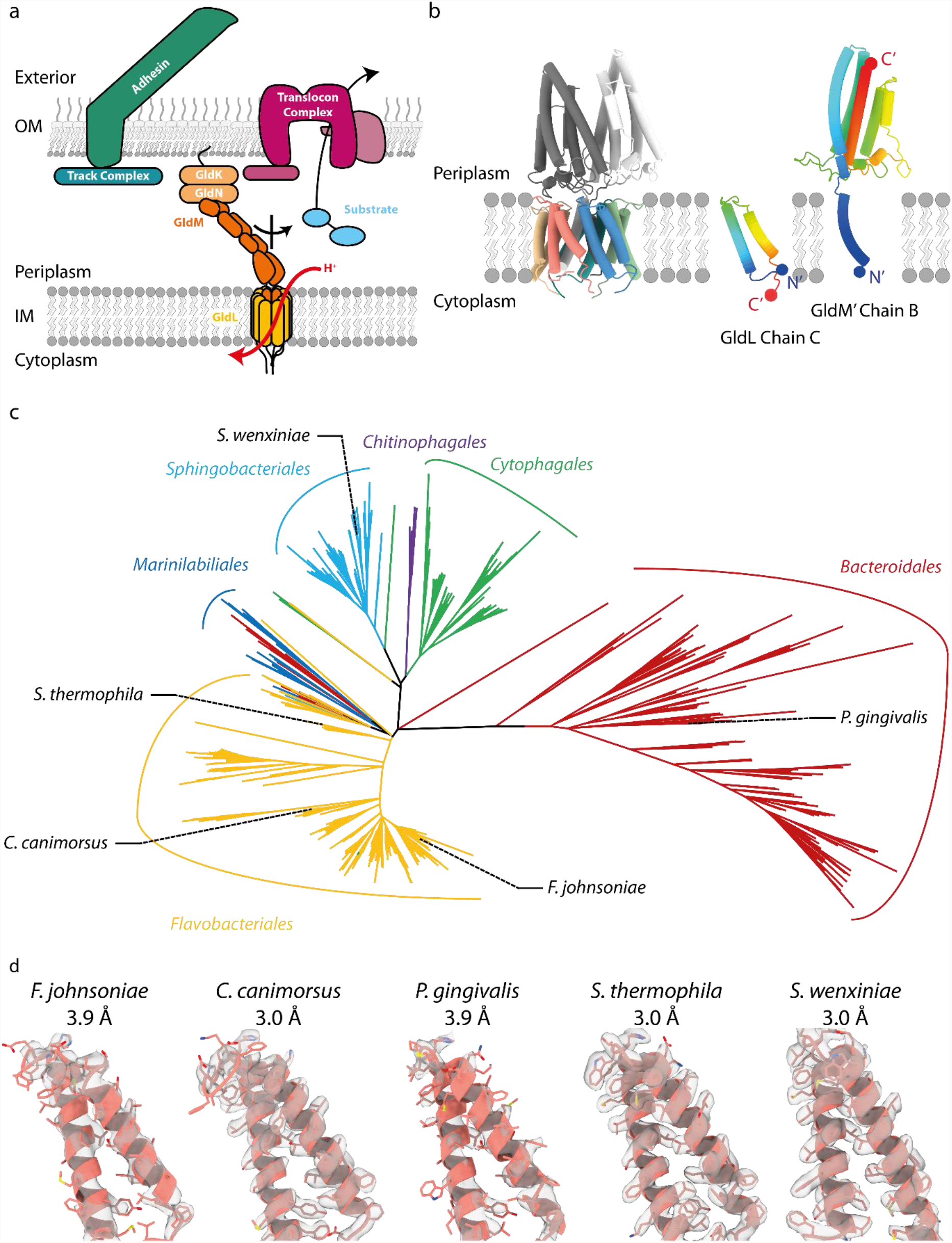
Role and phylogenetic diversity of the GldLM motor complex. **a** Cartoon illustrating the involvement of the GldLM motor complex in theT9SS and gliding motility. The GldLM motor converts electrochemical potential energy from the proton-motive force across the inner membrane (IM) into mechanical work which the periplasmic portion of GldM transfers across the periplasm to the outer membrane (OM). This mechanical energy is used to drive gliding adhesin movement (left) and protein transport through the T9SS (right). Coupling between these processes and GldM is thought to be mediated by a GldKN lipoprotein complex. **b** Cartoon representation of the structure of the *F. johnsoniae* GldLM′ complex solved previously ((11); PDB 6SY8, EMD-10893). (*Left*) Whole structure. The five GldL chains are colored salmon, blue, green, teal, and salmon and the two GldM chains are colored dark gray and white. (*Right*) Individual GldL and GldM chains are shown and rainbow colored from the N-terminus (blue) to the C-terminus (red). The most N-terminal (N′) and C-terminal (C′) modelled residues of each chain are marked with a sphere. **c** Maximum likelihood phylogenetic tree of GldL sequences in the *Bacteroidetes* phylum. Branches are colored by taxonomic order and the positions of proteins for which structures were determined are indicated. Note that the GldL sequence similarity matches the bacterial phylogenetic tree in most cases indicating that GldL has been predominantly vertically rather than horizontally transmitted. A phylogenetic tree for GldM is shown in Figure S1. **d** Increased resolution of the new T9SS/gliding motor complex structures shows improved side chain density. Chain GldL_c_ is shown for each species with EM density displayed at the same contour level.

The T9SS and the gliding motility apparatus share a motor complex that uses the proton-motive force (PMF) across the inner membrane to drive both protein transport and gliding adhesin movement at the outer membrane (11–13). The motor complex is formed from the integral inner membrane proteins GldL and GldM (11, 14). GldL has two transmembrane helices (TMHs) and a cytoplasmic domain. GldM has one TMH and a large periplasmic region which crystal structures have shown forms an extended dimer of four domains D1-D4 (11, 15, 16). The periplasmic region of GldM is long enough to span the periplasm to contact the outer membrane components of the T9SS and gliding motility apparatus.

We previously solved the structure of the core of the GldLM motor complex from the gliding bacterium *Flavobacterium johnsoniae* (11). This structure contains the full transmembrane region of the motor complex but the cytoplasmic domain of GldL was not visible and the periplasmic region of GldM had been genetically truncated after the first (D1) domain, forming a construct termed *Fjo*GldLM′ (11). The structure reveals that the T9SS/gliding motor is a GldL_5_GldM_2_ heteroheptamer in which the 10 GldL TMHs surround the two GldM TMHs (Figure 1b). The symmetry mismatch between the total number of GldL and GldM TMHs results in an inherently asymmetric relationship between the two types of subunit around the GldL ring. Amino acid substitutions showed that several conserved protonatable residues in the transmembrane helices of GldL and GldM are important for T9SS and gliding motility function. These residues are likely to be involved in coupling proton flow across the inner membrane to mechanical motions in the motor. Based on the organisation of the GldLM transmembrane helices, and the structural homology of the transmembrane part of GldLM to the ion-driven motor complexes that drive bacterial flagella (17, 18), we proposed that GldLM forms a rotary motor in which the GldM subunits rotate within the ring of GldL helices (11, 17). The periplasmic domain of GldM is then envisaged to transmit this rotary motion across the periplasm to the outer membrane components of the T9SS and gliding motility systems.

The structure of the *Fjo*GldLM′ complex was determined to a resolution (3.9 Å) at which only limited information can be inferred about the position of mechanistically important amino acid side chains. In addition, the protein was captured in a single conformational state providing only one snapshot of the catalytic mechanism of the enzyme. In this work we have sought to overcome these limitations in the structural characterization of the GldLM motor by determining structures of the motor complexes from a phylogenetically diverse range of organisms. We anticipated that some of these complexes would allow structure determination at improved resolution and in alternative conformational states. Here we present structures of the T9SS/gliding motor core from the human pathogens *P. gingivalis* and *Capnocytophaga canimorsus*, and the environmental bacteria *Schleiferia thermophila* and *Sphingobacterium wenxiniae*.

## Results and discussion

### The architecture of GldLM′ complexes is conserved across the *Bacteroidetes*

To structurally survey the diversity of gliding motility/T9SS motor complexes we selected proteins from a range of *Bacteroidetes* bacteria for recombinant expression in *Escherichia coli*. These proteins were chosen to maximise phylogenetic spread, to include proteins from organisms growing at a range of temperatures (psychrophiles to thermophiles) and from different environments (marine, fresh water, terrestrial, and commensals/pathogens), and to include examples both from gliding bacteria and from non-gliding bacteria with a T9SS. All constructs included the full length GldL homologue. For the GldM homologue we trialled constructs including different numbers of periplasmic domains. However, as with the earlier *F. johnsoniae* GldLM′ structure, we were only able to obtain structures by cryo-EM from complexes in which GldM was truncated after the first periplasmic D1 domain (GldM′), with the exception of one construct in which GldM was truncated after the D2 domain (GldM″). However, even in the latter case this effectively produced a GldLM′ structure as the D2 domain was not resolved, as discussed below.

Following screening for expression, purification, and cryo-freezing we were able to determine structures for the T9SS/gliding motor complexes of *P. gingivalis* (*Pgi*PorLM′), *Capnocytophaga canimorsus* (*Cca*GldLM″_peri_ and *Cca*GldLM″_TMH_), *Schleiferia thermophila* (*Sth*GldLM′), and *Sphingobacterium wenxiniae* (*Swe*GldLM′) (Figure 1, Table 1 and Figures S2-S7). *P. gingivalis* is a non-gliding oral pathogen, *C. canimorsus* is a dog commensal and opportunistic human pathogen, *S. thermophila* is a thermophile isolated from a hot spring (optimum growth temperature 50 °C), and *S. wenxiniae* was isolated from a waste water treatment plant. Note that in *P. gingivalis* the motor proteins are termed PorLM rather than GldLM. Figure 1c and Figure S1 show the phylogenetic positions of these motor proteins within the diversity of GldL and GldM proteins. The resolutions of the new structures ranged from 3.0 to 3.9 Å, with the higher-resolution structures allowing for more confident positioning of side chains than our previous 3.9 Å resolution *Fjo*GldLM′ structure (Figure 1d).

**Table 1.**
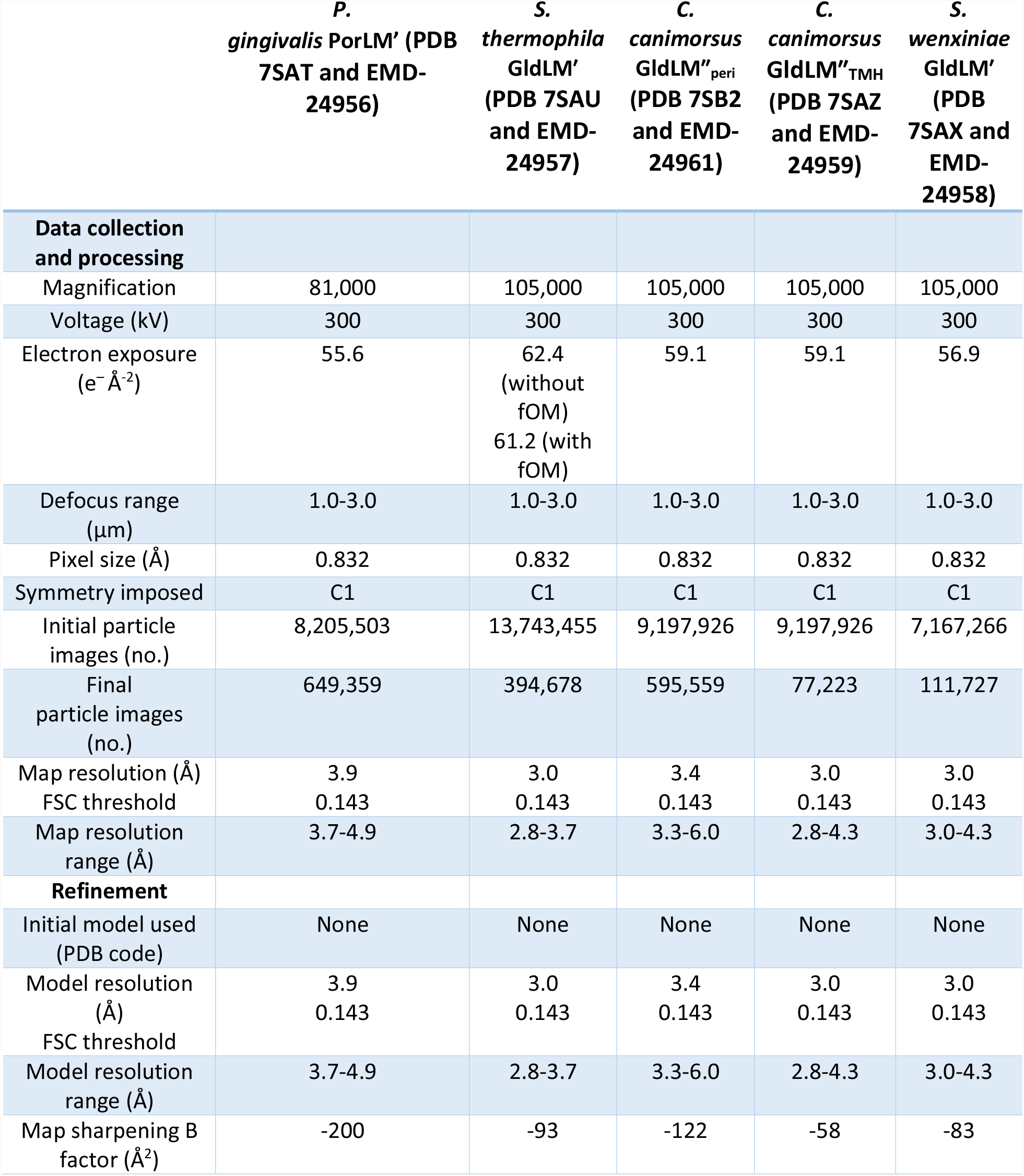
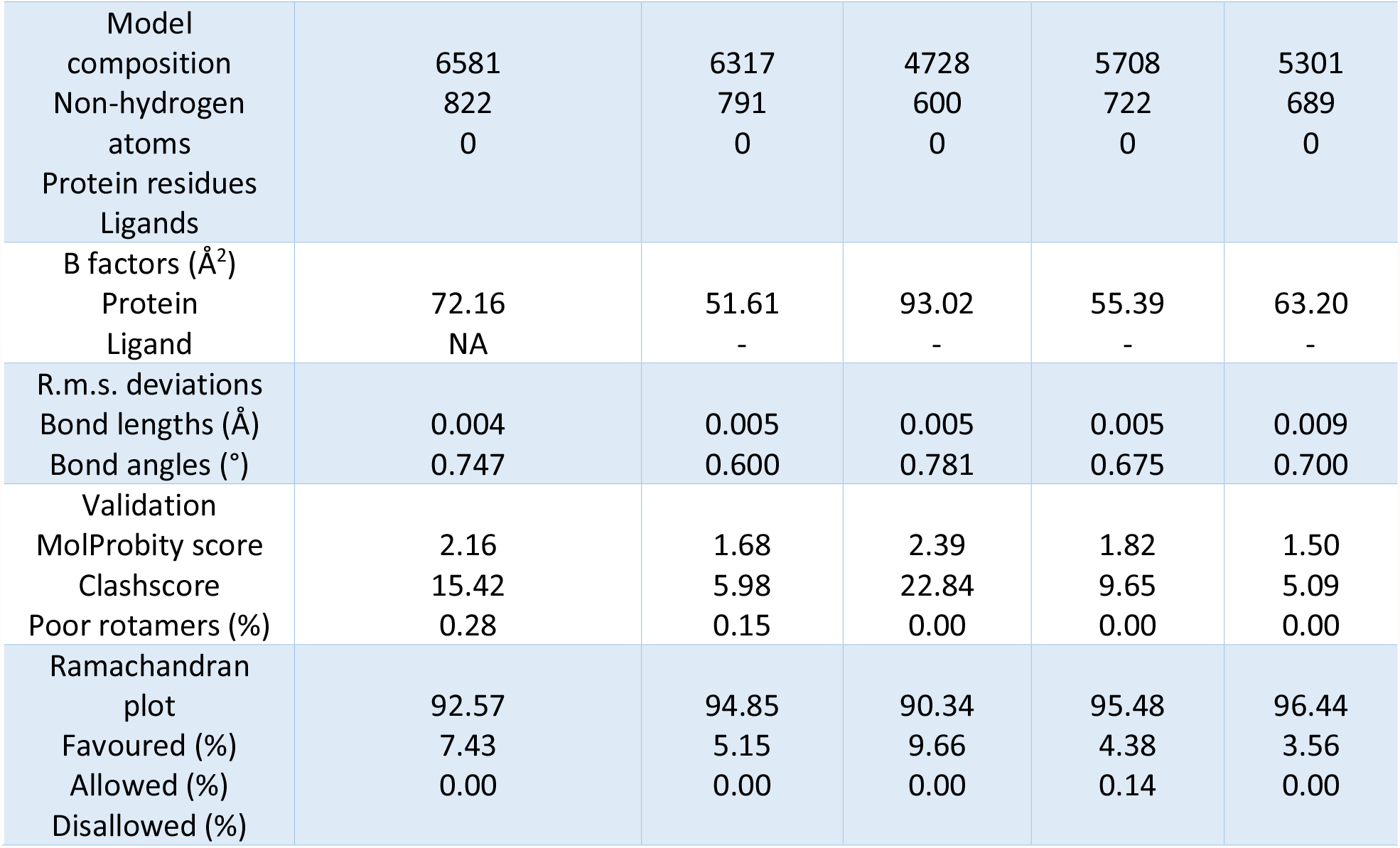
Cryo-EM data collection, refinement, and validation statistics for the *Pgi*PorLM′_C_, *Sth*GldLM′, *Cca*GldLM″_peri_, GldLM″_TMH_, and *Swe*GldLM′ structures.

The overall architecture of the *Fjo*GldLM′ complex is conserved in the four new motor complex structures, with five copies of GldL surrounding two copies of GldM (Figure 2a-c). As in the *Fjo*GldLM′ structure, only the trans-membrane helices and periplasmic loop of GldL were fully resolved, with almost all of the C-terminal cytoplasmic domain not seen in the structures (11). The periplasmic D1 domains of the GldM dimer were visible in all structures. The precise angles between the two copies of the D1 domain varied from 30° to 45° between structures. However, in all cases the D1 pairs adopted the splayed arrangement seen in the previous *Fjo*GldLM′ structure. This splayed arrangement contrasts with the closed arrangement of the D1 pair seen in crystal structures of the isolated periplasmic domain of GldM/PorM (15, 16), but is consistent with the low resolution cryo-EM structure of full length *Pgi*PorLM (11).

**Figure 2.**
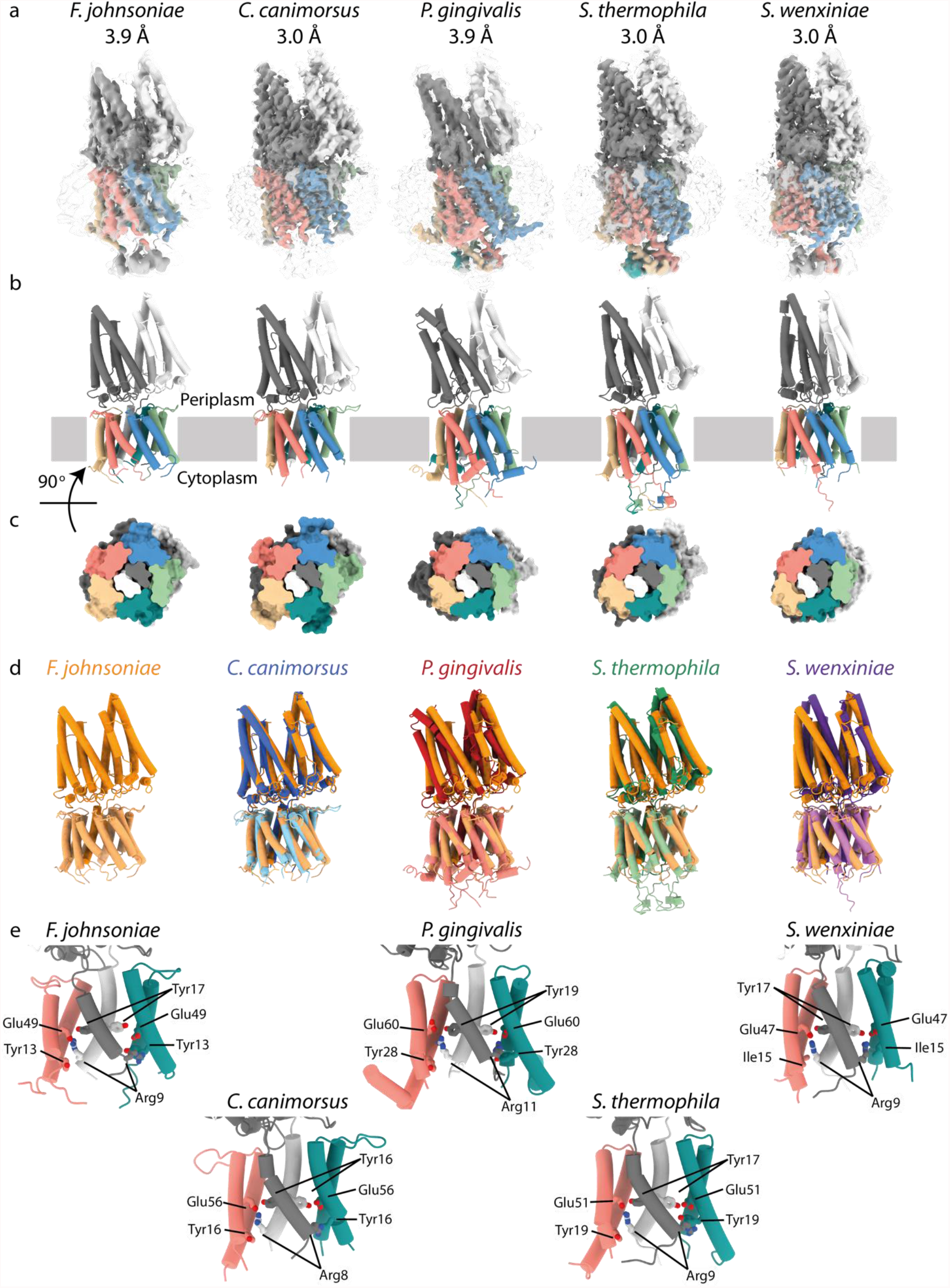
GldLM′ has conserved architecture across the *Bacteroidetes* phylum. **a** EM density maps for GldLM″_TMH_/GldLM′/PorLM′ complexes from the indicated species at high (colored by protein chain) and low (transparent) contour. The structure of *F. johnsoniae* GldLM′ was solved previously (11). The resolution of the structures is indicated above the panels. **b** Cartoon representations of the structures with chains colored as in **a. c** Slab through the protein density from **a** viewed from the cytoplasm and sliced approximately half-way through the membrane region. **d** The new GldLM″_TMH_/GldLM′/PorLM′ complex structures (colored as indicated) overlaid on *F. johnsoniae* GldLM′ (orange). GldM″_TMH_/GldM′/PorM′ subunits are shown in a darker shade than the GldL/PorL subunits. **e** Conservation of residues that are functionally important in *F. johnsoniae* GldLM (11) (top left hand panel) in other GldLM″_TMH_/GldLM/PorLM complexes. The proposed inter-subunit salt bridge is between the labelled Glu residue in the salmon GldL chain and the labelled Arg residue in the white GldM chain. For clarity chains GldL_D_, GldL_E_ and GldL_G_ are hidden for each structure.

Structural data were obtained from a *C. canimorsus* construct (*Cca*GldLM″) which includes both the D1 and D2 periplasmic domains of GldM. During 3D classification classes could be identified where either the D2 domain was well-resolved, or where the transmembrane portion of the complex was well-resolved (corresponding to the GldLM′ complex structures determined from other organisms), but never both (Figures S3, S4 & S8a-b). This suggests that the *Cca*GldLM″ complex is not able to adopt a conformation in which both the transmembrane helices and D2 domains are simultaneously ordered. In the *Cca*GldLM” structure with the ordered transmembrane helices (*Cca*GldLM″_TMH_), the D1 domains are splayed as in the other GldLM′/PorLM′ structures and in the low resolution structure of the complete PorLM complex (Figure S8 a,d). By contrast, in the *Cca*GldLM″ structure where the transmembrane domain is unresolved (*Cca*GldLM″_peri_), the D1 domains adopt a parallel orientation (Figure S8 b,c,e) similar to the isolated *Fjo*GldM periplasmic domain crystal structure (15) even though no crystal contacts are present. These observations suggest that splaying of the D1 domains is the most stable arrangement of the D1 domains in the intact motor complex, but leaves open the possibility that a parallel arrangement of the D1 domains might occur transiently during operation of the motor.

Mutagenesis has previously been used to identify residues within the transmembrane domain of the *F. johnsoniae* GldLM motor that are important for function (11). These residues are well-conserved in the new motor structures, with the exception that *Fjo*GldL Tyr13 is replaced by isoleucine in *S. wenxiniae* GldLM. The side chain positions of these residues can be assigned with more confidence due to the improved resolution of the *Cca*GldLM”_TMH_, *Sth*GldLM′, and *Swe*GldLM′ structures. Notably the orientation of these side chains is similar between the different motor structures where the resolution of the structures allows this judgement (Figure 2e). The side chain position is least well defined for the *Fjo*GldL Glu49 equivalent, which we previously proposed forms a salt bridge between one copy of GldL and Arg9 in one of the two *Fjo*GldM molecules (11). The poor definition of this residue is not surprising as glutamate sidechains are susceptible to damage by the electron beam during cryo-EM experiments, meaning that they are often not visible in EM density maps and cannot be accurately modelled (19). Nevertheless, in all the motor protein structures the side chain of the *Fjo*GldM Arg9 equivalent is oriented towards the *Fjo*GldL Glu49 equivalent suggesting that a salt bridge interaction between these residues is a conserved feature of the GldLM/PorLM complex.

### The membrane-proximal region of GldL forms a cage-like structure

The majority of the cytoplasmic region of GldL was not visible in any of our structures. However, the part that is immediately adjacent to the inner face of the cytoplasmic membrane was much better resolved in the *Pgi*PorLM′, *Sth*GldLM′, and *Swe*GldLM′ structures than in the previously determined *Fjo*GldLM′ structure. It can now be seen that extended coils from the C-terminal end of TMH2 form a cage-like structure below the detergent micelle (Figure 3a). In *Swe*GldLM′ the coil was only fully resolved for chain GldL_C_, which is involved in the putative salt bridge with chain GldM_B_. This coil is braced by the N-terminus of the adjacent chain GldL_D_, an interaction not seen in the other structures. The cage structure is best-resolved in the *Sth*GldLM′ complex, revealing that the constituent coils are held together by a network of hydrogen bonds and hydrophobic packing interactions between aspartate, tryptophan and tyrosine residues (Figure 3d). The cage structure exhibits high sequence conservation suggesting that it is of structural and/or functional importance (Figure 3d and Figure S9a). The cytoplasmic interactions between the GldL chains in the cage structure could help coordinate movements between subunits that do not contact each other in the TMH bundle. The surface of the cage is acidic (Figure S9b) creating a region of negative charge that may assist in the release of protons flowing through the transmembrane part of the motor complex. Alternatively, it may help maintain separation between the cytoplasmic domain of GldL and the negatively-charged phospholipid head groups of the cytoplasmic membrane.

**Figure 3.**
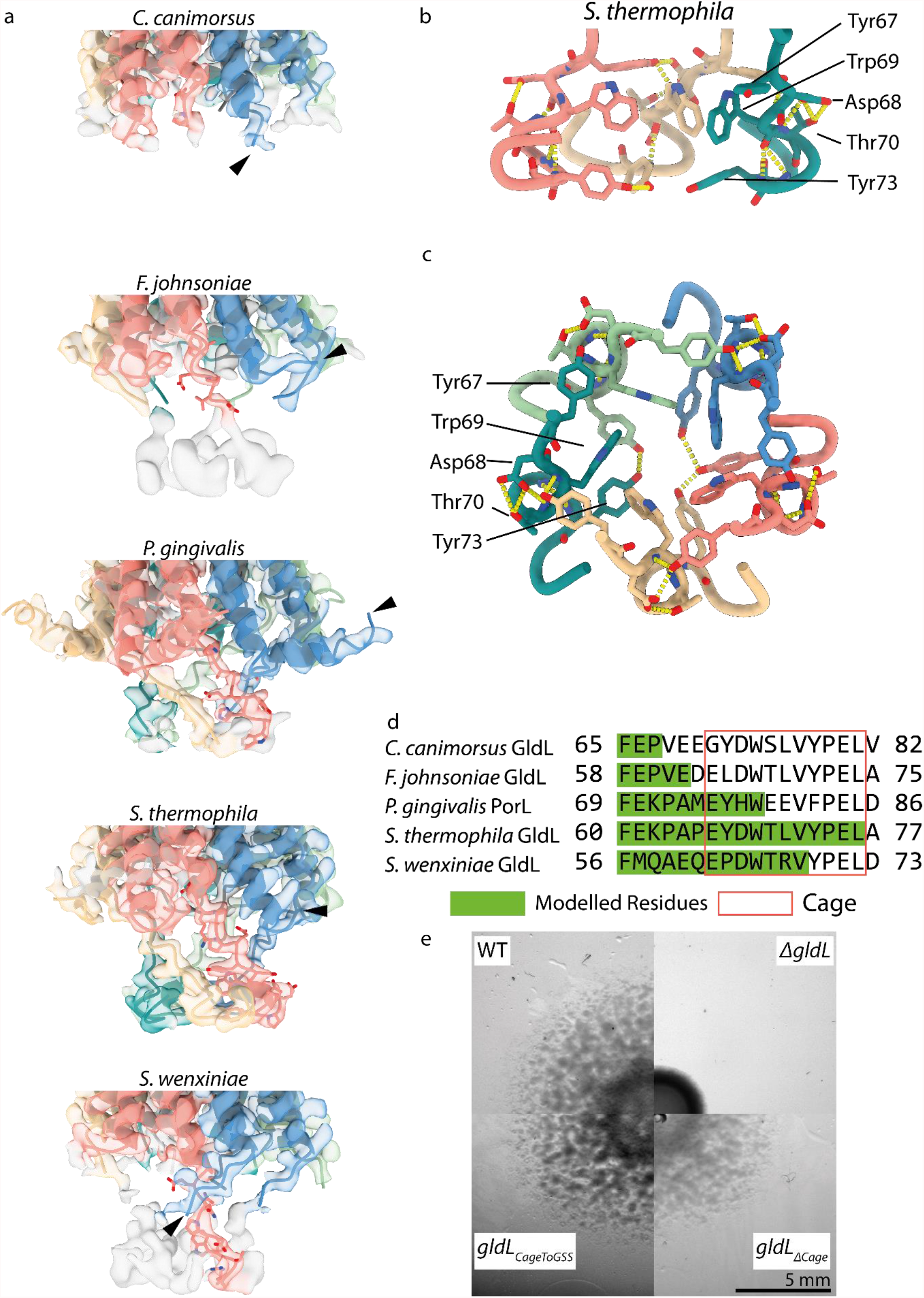
The membrane-proximal part of the GldL cytoplasmic domain forms a cage. **a** Overlay of EM density and the built model for the cytoplasmic region of each GldL structure. EM density is shown at the same contour level for all species. Side chains are shown for Chain C in the cage region. The most N-terminal residue modelled for Chain D is indicated with an arrowhead to highlight the bracing interaction between Chains C and D of *Swe*GldLM′. **b**,**c** Interaction network at the base of the cage-like structure in *Sth*GldLM′. The view direction is parallel to the membrane (**b**) or from within the TMH bundle (**c**). For clarity chains D and F are hidden in **b**. Hydrogen bonds are shown as yellow dashes. Selected side chains are displayed for the same residues in each chain and are labelled on Chain F (teal). For other residues, backbone atoms are shown if they form hydrogen bonds. **d** Sequence alignment of the cage region for the five GldL/PorL sequences. Residues which could be modelled for each structure are highlighted in green. Residues constituting the cage region are boxed in red. **e** Effects of modifications in the GldL cytoplasmic cage on *F. johnsoniae* gliding motility (spreading) on plates. The region of the *F. johnsoniae* sequence enclosed by the red box in **d** (residues E64-L74) was either substituted with GSSGSSGSSGS (*gldL*_*CagetoGSS*_) or deleted (*gldL*_*ΔCage*_). The results are representative of three independent experiments.

We investigated the importance of the cage structure to motor function through mutagenesis of the chromosomal *gldL* gene in the genetically tractable organism *F. johnsoniae*. Mutant cells in which the cage was completely deleted (removal of residues 64-74, *gldL*_*Δcage*_) had a small but reproducible gliding defect as measured by colony spreading on agar plates, whilst cells in which the cage sequence was replaced by a GSS repeat linker of the same length (*gldL*_*CageToGSS*_) showed no gliding defect (Figure 3e). The fact that changing the cage sequence to a GSS repeat had no effect on motility on agar indicates that the length of this cage region, rather than its precise sequence or structure, is most important to motor function.

### *P. gingivalis* PorL has a N-terminal helix that is absent in other structurally characterized GldL/PorL proteins

The *P. gingivalis* PorL protein has a N-terminal extension relative to the other motor proteins that we have structurally characterized (Figure 4a). This extension forms a helix that is not present in the other structures (Figure 4b). The helix points away from the trans-membrane GldL helix bundle at an angle of approximately 100° from TMH1 (Figure 4c,d) and approximately tangential to the circumference of the bundle (Figure 4b). The helix lies against the curved surface of the detergent micelle (Figure 4e) suggesting that *in vivo* it is likely to lie along the membrane surface with the positively charged N-terminus (Figure 4b,c) interacting with the negatively-charged phospholipid head groups. This helix may, therefore, play a role in stabilising the position of the PorLM complex in the cytoplasmic membrane. The presence of this N-terminal helix in *Pgi*PorLM′ and the bracing interaction seen between the N-terminus of GldL_D_ and the cytoplasmic region of GldL_C_ in the *Swe*GldLM′ structure noted in the last section suggests a role for the N-terminus of GldL in species-specific functional tuning of the motor complex.

**Figure 4.**
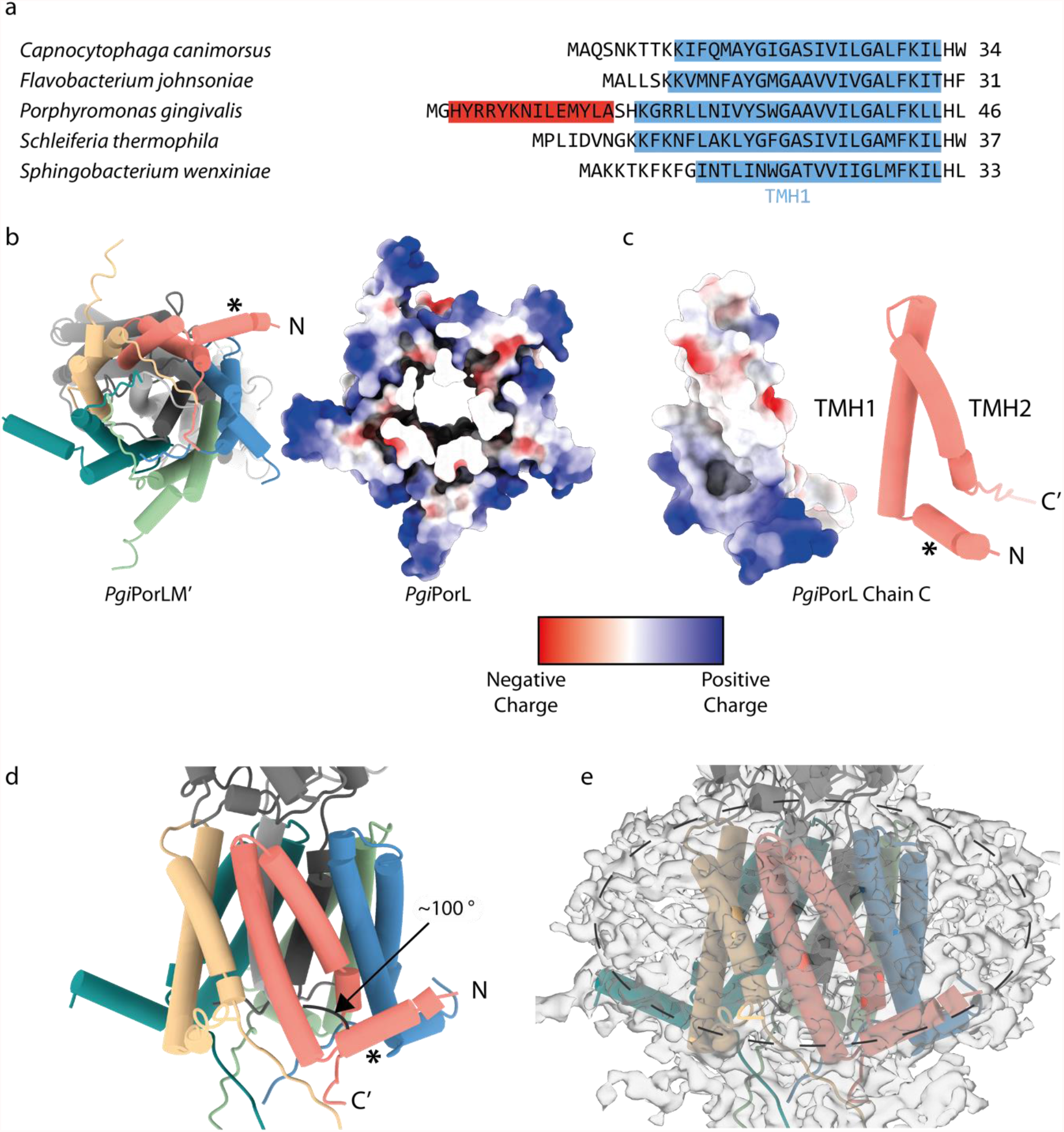
*P. gingivalis* PorL has a N-terminal membrane surface-associated helix. **a** Sequence alignment of the N-terminal regions of the structurally characterised GldL homologues. The first transmembrane helix of each sequence is highlighted in blue. The additional N-terminal helix of *Pgi*PorL is highlighted in red. **b** View from the cytoplasm of *P. gingivalis* PorLM′ in cartoon (left) and coulombic surface (right) representation. For clarity PorM′ is hidden in the coulombic view. **c** Coulombic surface (left) and cartoon (right) representations of *P. gingivalis* PorL Chain C viewed from within the membrane. **d** Side view of *Pgi*PorLM′ in cartoon representation. **e** Side view of the *Pgi*PorLM′ complex model overlaid with the EM density map displayed at low contour level. The approximate boundary of the detergent micelle is marked with a dashed line. In panels **b-d** * indicates the N-terminal helix and C′ indicates the most C-terminal modelled residue.

## Conclusion

This work, together with our previous study (11), provides the structures of the transmembrane cores of five T9SS/gliding motor complexes from species across three orders of the *Bacteroidetes*. These structures show that the architecture of the GldLM motor complex is well-conserved and imply that the mechanism by which the motor converts proton flow to mechanical movement is the same across the *Bacteroidetes*. The yield of the recombinant *S. wenxiniae* GldLM′ complex is much higher than that of the previously purified *Fjo*GldLM′ protein (Figure S2 and (11)) which should expedite future *in vitro* mechanistic studies of the T9SS/gliding motor. Future work should also explore how GldL and GldM interact with other components of the T9SS and gliding motility machinery to convert motor motions into the useful work of protein translocation and adhesin propulsion.

## Methods

### Bioinformatics analysis

A phylogenetic tree of GldL sequences was generated as follows. GldL sequences were obtained by BLAST searches against the UniRef90 database using the *F. johnsoniae* GldL, *P. gingivalis* PorL and *S. wenxiniae* GldL sequences as queries (20, 21). Each sequence in the UniRef90 database represents a cluster of sequences with more than 90% identity to the representative sequence. Searching against the UniRef90 database reduces the number of highly similar sequences in the results compared to searching against an unfiltered database. Sequences duplicated between the three BLAST searches, sequences representing clusters where the lowest common taxon was higher than order (and thus likely incorrectly phylogenetically assigned), and sequences from organisms outside the *Bacteroidetes* phylum were removed. The sequences were then aligned using Clustal Omega (22). A phylogenetic tree was inferred using the Maximum Likelihood function of program MEGA X with a JTT matrix-based model with default settings (23, 24). A phylogenetic tree of GldM sequences was generated using the same approach.

### Bacterial strains and growth conditions

Strains and plasmids used in this study are listed in Tables S1 & S2. For cloning procedures, *E. coli* cells were routinely grown in Luria Bertani medium (LB; (25)) at 37 °C with shaking. *F. johnsoniae* cells were routinely grown in casitone yeast extract (CYE) (26) medium at 30 °C with shaking. PY2 medium (27) was used to assess motility on agar plates. When required, kanamycin was added to LB medium at 30 µg ml^-1^ or to TB medium at 50 µg ml^-1^. When required, erythromycin was added at 100 µg ml^-1^.

### Genetic constructs

Primers used in this work are described in Table S3. All plasmid constructs were verified by sequencing.

GldL and C-terminally truncated and Twin-Strep-tagged GldM proteins (GldM′-TS/GldM″-TS) were expressed from vectors derived from the plasmid pT12 (28) under the control of rhamnose-inducible promoters.

Suicide vectors to genetically modify *F. johsoniae* were produced using the vectors pRHJ012 (11) and pYT354 (29), then introduced using into the *F. johnsoniae ΔgldL* strain Fl_082 (11) using *E. coli* strain S17-1 (30) as previously described (11).

Full details of all genetic constructs are given in the Supplemental Methods.

### Purification of protein complexes

Briefly, protein complexes were overexpressed in BL21(DE3) cells and then extracted from cell membranes using lauryl maltose neopentyl glycol (LMNG; Anatrace). Protein complexes were then affinity-purified using StrepTactin XT resin (IBA) and then further purified using size-exclusion chromatography. Full details of the purification scheme are given in the Supplemental Methods.

Typical yields per litre of cell culture were as follows: *Cca*GldLM″ 20 µg, *Pgi*PorLM′ 50 µg, *Sth*GldLM′ 6 µg, *Swe*GldLM′ 350 µg.

### Cryo-EM sample preparation and imaging

4 µl aliquots of purified samples at A_280_ = 1 were applied onto glow-discharged holey carbon-coated grids (Quantifoil 300 mesh, Au R1.2/1.3), then adsorbed for 10 s, blotted for 2 s at 100% humidity at 4 °C and plunge-frozen in liquid ethane using a Vitrobot Mark IV (FEI). To prepare samples with fluorinated octyl maltoside (fOM; Anatrace), proteins were concentrated to A_280_ = 3 and 13.5 µl was mixed with 1.5 µl of 7 mM fOM in Buffer W + 0.01% LMNG. All samples were centrifuged at 18,400*g* for 10 min at 4 °C immediately before grid preparation.

Data were collected using a Titan Krios G3 (FEI) operated at 300 kV fitted with either a GIF energy filter (Gatan) and a K2 Summit detector (Gatan) or a BioQuantum imaging filter (Gatan) and a K3 direct detection camera (Gatan). Full details of the data collection strategy are given in the Supplemental Methods.

### Cryo-EM data processing

Motion correction, dose weighting, contrast transfer function determination, particle picking, and initial particle extraction were performed using SIMPLE 3.0 (31). Gold-standard Fourier shell correlations (FSC) using the 0.143 criterion and local resolution estimations were calculated within RELION 3.1 (32).

In general, extracted particles were subjected to reference-free 2d classification in SIMPLE, followed by 3d classification in Relion using either the previously-solved *Fjo*GldLM′ map (11) or another map produced in this study as a reference. Classes with clear secondary structure detail were selected and used for 3d autorefinement. Successive rounds of Bayesian particle polishing, 3d classification and 3d autorefinement in Relion were used to generate the final maps for each dataset. A full description of the data processing strategy for each dataset is given in the Supplemental Methods.

### Model building and refinement

The Phyre^2^ server was used to generate homology models for each new sequence from the structure of *Fjo*GldLM′ (PDB: 6SY8) using one-to-one threading (33). These models were rigid body fit into the cryo-EM volume using Coot and residues were built *de novo* or removed as necessary in Coot (34). Rebuilding in globally-sharpened and local-resolution filtered maps was combined with real-space refinement in Phenix using secondary structure, rotamer and Ramachandran restraints to give the final models described in Table 1 (35, 36). Validation was done in Molprobity (37). Structures were analysed using ChimeraX (38), Pymol 2.3.3 (Schrodinger) and the Consurf server (39, 40).

### Measurement of gliding motility on agar

Strains were grown overnight in PY2 medium, washed once in PY2 medium, then resuspended in PY2 medium to an OD600 = 0.1. A 2-μl sample was then spotted onto PY2 agar plates. Plates were incubated at 25 °C for 48 h before imaging with a Zeiss AXIO Zoom MRm CCD camera and Zeiss software (ZenPro 2012, v.1.1.1.0).

## Acknowledgements

We thank K. Foster for providing additional imaging facilities and E. Furlong for preparing some cryo-EM grids

This work was supported by Wellcome Trust studentship 102164/Z/13/Z, Wellcome Trust Investigator Awards 107929/Z/15/Z and 219477/Z/19/Z, and ERC Advanced Award 833713. This research was supported in part by the Intramural Research Program of the NIH. We acknowledge the use of the Central Oxford Structural Microscopy and Imaging Centre (COSMIC). COSMIC was supported by a Wellcome Trust Collaborative Award 201536/Z/16/Z, the Wolfson Foundation, a Royal Society Wolfson Refurbishment Grant, the John Fell Fund, and the EPA and Cephalosporin Trusts.

## Author contributions

RHJ performed genetic and biochemical work. AH optimised protein purification with RHJ. JCD prepared cryo-EM grids and collected cryo-EM data. RHJ solved cryoEM structures and built models with advice from JCD. RHJ, JCD and SML analysed structures. RHJ, SML and BCB wrote the of the manuscript. All authors commented on the manuscript and approved the final version.

## Data availability

The cryo-EM volumes and atomic coordinates presented in this paper have been deposited in the Electron Microscopy Data Bank (EMDB) and Protein Data Bank (PDB), respectively, with the following accession codes: *Cca*GldLM″_TMH_ -PDB 7SAZ and EMD-24959, *Cca*GldLM″_peri_ -PDB 7SB2 and EMD-24961, *Pgi*PorLM′ - PDB 7SAT and EMD-24956, *Sth*GldLM′ - PDB 7SAU and EMD-24957, and *Swe*GldLM′ - PDB 7SAX and EMD-24958.

## Supplemental Material Legends

**Table S1. Bacterial strains used in this study**.

**Table S2. Plasmids used in this study**

**Table S3. Oligonucleotides used in this study**.

**Figure S1 Maximum likelihood phylogenetic tree of GldM sequences in the *Bacteroidetes* phylum**. Branches are colored by taxonomic order and the positions of proteins for which structures were determined are indicated.

**Figure S2 Purification of GldLM′′, PorLM′, and GldLM′ complexes. a** Size exclusion-chromatography traces for the indicated protein complexes. Proteins were analysed using a Superose 6 10/300 Increase column (Cytiva). The peaks used to prepare cryo-EM grids are indicated with arrowheads. **b-e** SDS-PAGE gels for the fractions used to make cryo-EM grids for the indicated protein complexes.

**Figure S3 Data processing workflow for *Cca*GldLM″**_**peri**_ **map. a** Example micrograph for the *Cca*GldLM′′ sample collected at approximately 2.6 µm defocus. Scale bar 500 Å. **b** Representative 2D class averages used to produce the initial model. Scale bar 100 Å. **c** Representative 2D class averages used to produce the *Cca*GldLM″_peri_ map. Scale bar 100 Å. **d** Data processing workflow for the *Cca*GldLM″_peri_ map. The handedness of the maps was flipped between the left and right columns once helices were visible. **e** Local resolution estimates (in Å) for the sharpened *Cca*GldLM″_peri_ map. **f** Fourier Shell Correlation (FSC) plot for the *Cca*GldLM″_peri_ map. The resolution at the gold-standard cut-off (FSC = 0.143) is indicated by the dashed line. Curves: Red, phase-randomised; Green, unmasked; Blue, masked; black, MTF-corrected.

**Figure S4 Data processing workflow for *Cca*GldLM″**_**TMH**_ **map. a** Representative 2D class averages used to produce the *Cca*GldLM″_TMH_ map. Scale bar 100 Å. **b** Data processing workflow for the *Cca*GldLM″_TMH_ map. **c** Local resolution estimates (in Å) for the sharpened *Cca*GldLM″_TMH_ map. **d** FSC plot for the *Cca*GldLM″_TMH_ map. The resolution at the gold-standard cut-off (FSC = 0.143) is indicated by the dashed line. Curves: Red, phase-randomised; Green, unmasked; Blue, masked; black, MTF-corrected.

**Figure S5 Data processing workflow for *Pgi*PorLM′ map. a** Example micrograph for the initial *Pgi*PorLM′ dataset collected with a K2 detector at approximately 1.8 µm defocus. Scale bar 500 Å. **b** Representative 2D class averages from the K2 dataset used for initial processing. Scale bar 100 Å. **c** Example micrograph from the second *Pgi*PorLM′ dataset collected with a K3 detector at approximately 1.8 µm defocus. Scale bar 500 Å. **d** Representative 2D class averages from the K3 dataset used for final processing. Scale bar 100 Å. **e** Data processing workflow for the *Pgi*PorLM′ map. Scale bar on 2D class averages 100 Å. **f** Local resolution estimates (in Å) for the sharpened *Pgi*PorLM′ map. **g** Fourier Shell Correlation (FSC) plot for the *Pgi*PorLM′ map. The resolution at the gold-standard cut-off (FSC = 0.143) is indicated by the dashed line. Curves: Red, phase-randomised; Green, unmasked; Blue, masked; black, MTF-corrected.

**Figure S6 Data processing workflow for *Sth*GldLM′ map. a** Example micrograph for the initial *Sth*GldLM′ dataset collected with a K2 detector at a defocus of approximately 2.7 µm. **b** Example micrograph for the *Sth*GldLM′ + fOM dataset collected with a K3 detector at a defocus of approximately 1.7 µm. **c** Example micrograph for the *Sth*GldLM′ dataset collected with a K3 detector at defocus of approximately 1.7 µm. **d** Representative 2D class averages for the K2 dataset used in initial processing. **e** Data processing workflow for the initial dataset collected with a K2 detector. **f** Representative 2D class averages for the *Sth*GldLM′ + fOM (left) and *Sth*GldLM′ only (right) K3 datasets. **g** Data processing workflow for the combined datasets collected with a K3 detector. In the bottom-left panel each cylinder represents a view orientation, and the height of the cylinder corresponds to the number of views of that orientation. **h** Local resolution estimates (in Å) for the sharpened *Sth*GldLM′ map. **i** FSC plot for the *Sth*GldLM′ map. The resolution at the gold-standard cut-off (FSC = 0.143) is indicated by the dashed line. Curves: Red, phase-randomised; Green, unmasked; Blue, masked; black, MTF-corrected. Scale bars for micrographs 500 Å. Scale bars for 2D class averages 100 Å.

**Figure S7 Data processing workflow for *Swe*GldLM′ map. a** Example micrograph for *Swe*GldLM′ dataset collected with K3 detector at defocus of approximately 1.5 µm. Scale bar 500 Å. **b** Representative 2D class averages for initial particle selection. Scale bar 100 Å. **c** Data processing workflow for initial particle selection. **d** Representative 2D class averages for second, side-view focused particle selection. Scale bar 100 Å. **e** Data processing workflow for second particle selection. **f** Local resolution estimates (in Å) for the sharpened *Swe*GldLM′ map. **g** Fourier Shell Correlation (FSC) plot for the *Swe*GldLM′ map. The resolution at the gold-standard cut-off (FSC = 0.143) is indicated by the dashed line. Curves: Red, phase-randomised; Green, unmasked; Blue, masked; black, MTF-corrected.

**Figure S8 Comparison of *Cca*GldLM″**_**TMH**_ **and *Cca*GldLM″**_**peri**_ **structures. a** *Cca*GldLM″_TMH_ EM density map. GldL is colored light blue and GldM″_TMH_ is colored dark blue. **b** *Cca*GldLM″_**peri**_ EM density map. **c** Protein model of *Cca*GldLM″_peri_. It was not possible to build protein into the transmembrane density. **d** Overlay of the *Cca*GldLM″_TMH_ and *Cca*GldLM″_peri_ structures showing the different relative orientations of the GldM D1 domains. Proteins are colored as in (a) and (c). **e** Overlay of *Cca*GldLM″_peri_ (yellow) and *Fjo*GldM_peri_ (PDB 6ey4)(orange) structures, showing the similarity in GldM D1 domain orientations. All maps and models in this figure were aligned to the second helix of *Cca*GldLM″_peri_ Chain A, indicated with a dashed line in (c).

**Figure S9 Conservation and charge analysis for the cage region of *Sth*GldLM′. a** Sequence conservation analysis of the cage region of *S. thermophila* GldL using the program Consurf (10, 11). **b**,**c** Coulombic potential representation for the cage region of *S. thermophila* GldL. The view direction is parallel to the membrane (**b**) or from within the TMH bundle (**c**).For clarity, the first TMH and N-terminal residues are hidden.

## References

1. Sato K, Naito M, Yukitake H, Hirakawa H, Shoji M, McBride MJ, Rhodes RG, Nakayama K. 2010. A protein secretion system linked to Bacteroidete gliding motility and pathogenesis. Proc Natl Acad Sci 107:276–81.

2. McBride MJ. 2019. Bacteroidetes Gliding Motility and the Type IX Secretion System. Microbiol Spectr 7:PSIB-0002-2018.

3. Lauber F, Deme JC, Lea SM, Berks BC. 2018. Type 9 secretion system structures reveal a new protein transport mechanism. Nature 564:77–82.

4. Lasica AM, Goulas T, Mizgalska D, Zhou X, De Diego I, Ksiazek M, Madej M, Guo Y, Guevara T, Nowak M, Potempa B, Goel A, Sztukowska M, Prabhakar AT, Bzowska M, Widziolek M, Thøgersen IB, Enghild JJ, Simonian M, Kulczyk AW, Nguyen KA, Potempa J, Gomis-Rüth FX. 2016. Structural and functional probing of PorZ, an essential bacterial surface component of the type-IX secretion system of human oral-microbiomic Porphyromonas gingivalis. Sci Rep 6:37708.

5. Nakayama K. 2015. Porphyromonas gingivalis and related bacteria: from colonial pigmentation to the type IX secretion system and gliding motility. J Periodontal Res 50:1–8.

6. Guo Y, Hu D, Guo J, Wang T, Xiao Y, Wang X, Li S, Liu M, Li Z, Bi D, Zhou Z. 2017. Riemerella anatipestifer Type IX secretion system is required for virulence and gelatinase secretion. Front Microbiol 8:2553.

7. Yuan H, Huang L, Wang M, Jia R, Chen S, Liu M, Zhao X, Yang Q, Wu Y, Zhang S, Liu Y, Zhang L, Yu Y, You Y, Chen X, Zhu D, Cheng A. 2019. Role of the gldK gene in the virulence of Riemerella anatipestifer. Poult Sci https://doi.org/10.3382/ps/pez028.

8. Li N, Zhu Y, LaFrentz BR, Evenhuis JP, Hunnicutt DW, Conrad RA, Barbier P, Gullstrand CW, Roets JE, Powers JL, Kulkarni SS, Erbes DH, García JC, Nie P, McBride MJ. 2017. The Type IX Secretion System Is Required for Virulence of the Fish Pathogen Flavobacterium columnare. Appl Environ Microbiol 83:e01769–17.

9. Cabral L, Persinoti GF, Paixao DAA, Martins MP, Chinaglia M, Morais MAB, Pirolla RAS, Generoso WC, Maciel LF, Lombard V, Henrissat B, Murakami MT. 2021. Gut microbiome of capybara, the Amazon master of the grasses, harbors unprecedented enzymatic strategies for plant glycans breakdown. Prepr Version 1 Available Res Sq https://doi.org/10.21203/rs.3.rs-456076/v1.

10. Naas AE, Solden LM, Norbeck AD, Brewer H, Hagen LH, Heggenes IM, McHardy AC, Mackie RI, Paša-Tolić L, Arntzen MØ, Eijsink VGH, Koropatkin NM, Hess M, Wrighton KC, Pope PB. 2018. “Candidatus Paraporphyromonas polyenzymogenes” encodes multi-modular cellulases linked to the type IX secretion system. Microbiome 6:44.

11. Hennell James R, Deme JC, Kjær A, Alcock F, Silale A, Lauber F, Johnson S, Berks BC, Lea SM. 2021. Structure and mechanism of the proton-driven motor that powers type 9 secretion and gliding motility. Nat Microbiol 6:221–223.

12. Dzink-Fox JAL, Leadbetter ER, Godchaux W. 1997. Acetate acts as a protonophore and differentially affects bead movement and cell migration of the gliding bacterium Cytophaga johnsonae (Flavobacterium johnsoniae). Microbiology 143:3693–3701.

13. Nakane D, Sato K, Wada H, McBride MJ, Nakayama K. 2013. Helical flow of surface protein required for bacterial gliding motility. Proc Natl Acad Sci 110:11145–50.

14. Vincent MS, Canestrari MJ, Leone P, Stathopulos J, Ize B, Zoued A, Cambillau C, Kellenberger C, Roussel A, Cascales E. 2017. Characterization of the Porphyromonas gingivalis Type IX Secretion Trans-envelope PorKLMNP Core Complex. J Biol Chem 292:3252–3261.

15. Leone P, Roche J, Vincent MS, Tran QH, Desmyter A, Cascales E, Kellenberger C, Cambillau C, Roussel A. 2018. Type IX secretion system PorM and gliding machinery GldM form arches spanning the periplasmic space. Nat Commun 9:429.

16. Sato K, Okada K, Nakayama K, Imada K. 2020. PorM, a core component of bacterial type IX secretion system, forms a dimer with a unique kinked-rod shape. Biochem Biophys Res Commun 532:114–119.

17. Deme JC, Johnson S, Vickery O, Aron A, Monkhouse H, Griffiths T, James RH, Berks BC, Coulton JW, Stansfeld PJ, Lea SM. 2020. Structures of the stator complex that drives rotation of the bacterial flagellum. Nat Microbiol 5:1553–1564.

18. Santiveri M, Roa-Eguiara A, Kühne C, Wadhwa N, Hu H, Berg HC, Erhardt M, Taylor NMI. 2020. Structure and Function of Stator Units of the Bacterial Flagellar Motor. Cell 183:244–257.

19. Hattne J, Shi D, Glynn C, Zee C-T, Gallagher-Jones M, Martynowycz MW, Rodriguez JA, Gonen T. 2018. Analysis of Global and Site-Specific Radiation Damage in Cryo-EM. Structure 26:759–766.

20. Altschul SF, Miller W, Gish W, Myers EW, Lipman DJ. 1990. Basic Local Alignment Search Tool. J Mol Biol 215:403–410.

21. Suzek BE, Wang Y, Huang H, McGarvey PB, Wu CH. 2015. UniRef clusters: A comprehensive and scalable alternative for improving sequence similarity searches. Bioinformatics 31:926–932.

22. Madeira F, Park YM, Lee J, Buso N, Gur T, Madhusoodanan N, Basutkar P, Tivey ARN, Potter SC, Finn RD, Lopez R. 2019. The EMBL-EBI search and sequence analysis tools APIs in 2019. Nucleic Acids Res 47:W636–W641.

23. Kumar S, Stecher G, Li M, Knyaz C, Tamura K. 2018. MEGA X: Molecular evolutionary genetics analysis across computing platforms. Mol Biol Evol 35:1547–1549.

24. Jones DT, Taylor WR, Thornton JM. 1992. The rapid generation of mutation data matrices from protein sequences. Bioinformatics 8:275–282.

25. Miller JH. 1972. Experiments in molecular genetics. Cold Spring Harbor Laboratory, Cold Spring Harbor, New York.

26. McBride MJ, Kempf MJ. 1996. Development of techniques for the genetic manipulation of the gliding bacterium Cytophaga johnsonae. J Bacteriol 178:583–590.

27. Agarwal S, Hunnicutt DW, McBride MJ. 1997. Cloning and characterization of the Flavobacterium johnsoniae (Cytophaga johnsonae) gliding motility gene, gldA. Proc Natl Acad Sci 94:12139–12144.

28. Dietsche T, Tesfazgi Mebrhatu M, Brunner MJ, Abrusci P, Yan J, Franz-Wachtel M, Schärfe C, Zilkenat S, Grin I, Galán JE, Kohlbacher O, Lea S, Macek B, Marlovits TC, Robinson CV, Wagner S. 2016. Structural and Functional Characterization of the Bacterial Type III Secretion Export Apparatus. PLOS Pathog 12:e1006071.

29. Zhu Y, Thomas F, Larocque R, Li N, Duffieux D, Cladière L, Souchaud F, Michel G, McBride MJ. 2017. Genetic analyses unravel the crucial role of a horizontally acquired alginate lyase for brown algal biomass degradation by Zobellia galactanivorans. Environ Microbiol 19:2164–2181.

30. Simon R, Priefer U, Pühler A. 1983. A broad host range mobilization system for in vivo genetic engineering: Transposon mutagenesis in Gram-negative bacteria. Bio/Technology 1:784–791.

31. Caesar J, Reboul CF, Machello C, Kiesewetter S, Tang ML, Deme JC, Johnson S, Elmlund D, Lea SM, Elmlund H. 2020. SIMPLE 3.0. Stream single-particle cryo-EM analysis in real time. J Struct Biol X 4:100040.

32. Zivanov J, Nakane T, Forsberg BO, Kimanius D, Hagen WJH, Lindahl E, Scheres SHW. 2018. New tools for automated high-resolution cryo-EM structure determination in RELION-3. eLife 7:e42166.

33. Kelley LA, Mezulis S, Yates CM, Wass MN, Sternberg MJE. 2015. The Phyre2 web portal for protein modeling, prediction and analysis. Nat Protoc 10:845–858.

34. Emsley P, Lohkamp B, Scott WG, Cowtan KD. 2010. Features and development of Coot. Acta Crystallogr D66:486–501.

35. Afonine PV, Poon BK, Read RJ, Sobolev OV, Terwilliger TC, Urzhumtsev A, Adams PD. 2018. Real-space refinement in PHENIX for cryo-EM and crystallography. Acta Crystallogr D74:531–544.

36. Liebschner D, Afonine PV, Baker ML, Bunkoczi G, Chen VB, Croll TI, Hintze B, Hung LW, Jain S, McCoy AJ, Moriarty NW, Oeffner RD, Poon BK, Prisant MG, Read RJ, Richardson JS, Richardson DC, Sammito MD, Sobolev OV, Stockwell DH, Terwilliger TC, Urzhumtsev AG, Videau LL, Williams CJ, Adams PD. 2019. Macromolecular structure determination using X-rays, neutrons and electrons: Recent developments in Phenix. Acta Crystallogr D75:861–877.

37. Williams CJ, Headd JJ, Moriarty NW, Prisant MG, Videau LL, Deis LN, Verma V, Keedy DA, Hintze BJ, Chen VB, Jain S, Lewis SM, Arendall WB, Snoeyink J, Adams PD, Lovell SC, Richardson JS, Richardson DC. 2018. MolProbity: More and better reference data for improved all-atom structure validation. Protein Sci 27:293–315.

38. Pettersen EF, Goddard TD, Huang CC, Meng EC, Couch GS, Croll TI, Morris JH, Ferrin TE. 2021. UCSF ChimeraX: Structure visualization for researchers, educators, and developers. Protein Sci 30:70–82.

39. Ashkenazy H, Abadi S, Martz E, Chay O, Mayrose I, Pupko T, Ben-Tal N. 2016. ConSurf 2016: an improved methodology to estimate and visualize evolutionary conservation in macromolecules. Nucleic Acids Res 44:W344–W350.

40. Landau M, Mayrose I, Rosenberg Y, Glaser F, Martz E, Pupko T, Ben-Tal N. 2005. ConSurf 2005: The projection of evolutionary conservation scores of residues on protein structures. Nucleic Acids Res 33:W299–W302.

