## Supplemental Information for "Structures of the Type IX Secretion/gliding motility motor from across the phylum *Bacteroidetes*"

#### Supplemental Methods

##### Genetic constructs

Strains and plasmids used in this study are listed in Tables S1 & S2. Primers used in this work are described in Table S3. All plasmid constructs were verified by sequencing.

Gene synthesis (Invitrogen GeneArt) was used to create plasmid pMA-RQ_SthPKLMN containing genes RS08900 (*sprF* homologue), RS08905 (*gldK*), RS08910 (*gldL*), RS08915 (*gldM*), and RS08920 (*gldN*) from *S. thermophila* str Yellowstone with a Twin-Strep tag coding sequence added to the end of *gldL*. Quikchange (Agilent) mutagenesis was used with primers RHJ035-040 to remove BamHI, NcoI and NdeI restriction sites by silent mutation, yielding plasmid pRHJ113. A vector for the co-expression of *S. thermophila* GldL and C-terminally truncated and Twin-Strep-tagged GldM (GldM'-TS) under the control of a rhamnose-inducible promoter was produced as follows. The fragment encoding GldL was amplified using primers RHJ177 and RHJ178. The intergenic region and a fragment encoding the first 229 amino acids of GldM were amplified using primers RHJ179 and RHJ203. These fragments were assembled by Gibson cloning with plasmid pT12 (1) linearised using primers RHJ162 and RHJ163 to give plasmid pRHJ117.

A vector for the co-expression of *P. gingivalis* PorL and C-terminally truncated and Twin-Strep-tagged PorM (PorM'-TS) under the control of a rhamnose-inducible promoter was produced as follows. The chromosomal region encoding PorL and the first 227 amino acids of PorM was amplified from *P. gingivalis* ATCC 33277 genomic DNA using primers RHJ168 and RHJ202. The plasmid pT12 was linearised using primers RHJ162 and RHJ163. The two fragments were then assembled by Gibson cloning to give plasmid pRHJ118. An analogous strategy using the primers listed in Table S3 was used to create plasmids pRHJ170 and pRHJ174 expressing *S. wenxiniae* DSM 22789 GldLM'-TS and *C. canimorsus* str. Cc5 GldLM"-TS motor complexes, respectively.

A suicide vector to delete codons E64 to L74 of *gldL* was produced as follows. pRHJ012 (2), containing a 5.3 kb region including *gldL* and the surrounding chromosomal regions was linearised by amplification with primers RHJ617 and RHJ618, which introduce the desired deletion. The resulting fragment was re-circularised by Gibson cloning to give pRHJ237. The fragment containing the mutated *gldL* sequence and adjacent regions was then amplified from pRHJ237 with primers RHJ341 and RHJ342. The vector pYT354 (3) was linearised by digestion with BamHI and SalI. The two fragments were then assembled by Gibson cloning to give plasmid pRHJ240. An analogous strategy was used to produce pRHJ241 where codons E64 to L74 of *gldL* are replaced with a GSSGSSGSSGS coding sequence, using the primers described in Table S3.

Suicide plasmids were introduced into *F. johnsoniae* strains by biparental mating using *E. coli* S17-1 (4) as the donor strain and *ΔgldL* strain Fl_082 (2) as the recipient, as previously described (5). Erythromycin was used to select cells with a chromosomally integrated suicide plasmid. One of the resulting clones was grown overnight in CYE without antibiotic to allow for loss of the plasmid backbone, and then plated onto CYE agar containing 5% sucrose. Sucrose-resistant colonies were screened by PCR for the presence of the desired chromosomal modification and then verified by sequencing.

#### Purification of protein complexes

*Pgi*PorLM′, *Swe*GldLM′, and *Cca*GldLM″ complexes were overproduced from plasmids pRHJ118, pRHJ170, and pRHJ174, respectively, as follows. Colonies of BL21(DE3) cells carrying the appropriate plasmid were used to inoculate 50 ml cultures of TB medium (6) with kanamycin and grown at 37 °C with shaking for 6-8 h. Cells were diluted to optical density at 600 nm ($OD_{600}$) of 0.02 in TB medium with kanamycin and 0.1% L-rhamnose then grown at 37 °C with shaking for 13 h. Cells were harvested by centrifugation at 5,000*g* for 15 min at 4 °C. Cells were washed once in Dulbecco A phosphate-buffered saline (Gibco) and stored at -20 °C until further use.

All purification steps were carried out at 4 °C except as otherwise noted. The frozen cell pellet was thawed at room temperature and then resuspended using a Dounce tissue grinder on ice in 4 ml per gram of cells of Buffer W (100 mM Tris-HCl pH 8.0, 150 mM NaCl, 1 mM EDTA) supplemented with 30 μg $ml^{-1}$ DNase I, 100 μg $ml^{-1}$ lysozyme, and 1 tablet per 100 ml SIGMA*FAST* (Merck) protease inhibitor cocktail. The cells were then disrupted using an Emulsiflex homogeniser operated at 100 MPa. The cell lysate was centrifuged at 24,000*g* for 30 min to remove debris and then centrifuged at 210,000*g* for 90 min to harvest cell membranes. Membranes were resuspended in 8 ml Buffer W per g membranes then 1 ml of 10% (w/v) lauryl maltose neopentyl glycol (LMNG; Anatrace) was added per gram of membrane and the suspension was gently stirred for 2 h. Unsolubilised material was removed by centrifugation at 100,000*g* for 30 min. The resulting supernatant was passed through a 5-ml StrepTactin XT cartridge (IBA). The column was washed with 10 column volumes of Buffer W with 0.02% LMNG then protein was eluted in 20 1 ml fractions of Buffer W with 0.01% LMNG and 50 mM D-biotin (IBA). Fractions containing GldL/PorL and GldM′/PorM′ were identified by $A_{280}$ and SDS-PAGE, concentrated to 500 µl using a 100-kDa molecular weight cut-off (MWCO) Amicon Ultra-15 centrifugal filter, then injected onto a Superose 6 10/300 Increase GL size-exclusion chromatography column (GE Healthcare) equilibrated in Buffer W + 0.01% LMNG at room temperature. Fractions containing purified complexes were identified by SDS-PAGE and concentrated using a GE Healthcare 100-kDa MWCO Vivaspin 500 concentrator. Protein concentrations were determined spectrophotometrically assuming that an $A_{280}$ of 1 is equivalent to 1 mg $ml^{-1}$.

*Sth*GldLM′ complexes were produced from pRHJ117 as above except that all steps after cell disruption were carried out at 15 °C (centrifugation steps) or at room temperature.

#### Cryo-EM sample preparation and imaging

Initial data for *Pgi*PorLM′ and *Sth*GldLM′ were collected in counting mode on a Titan Krios G3 (FEI) operating at 300 kV with a GIF energy filter (Gatan) and K2 Summit detector (Gatan) at 165,000x magnification using a pixel size of 0.822 Å and a total dose of 48 $e^{-}$ $Å^{-2}$ over 32 fractions.

The final *Pgi*PorLM′ dataset was collected in counted super-resolution mode on a Titan Krios G3 (FEI) operating at 300 kV with a BioQuantum imaging filter (Gatan) and K3 direct detection camera (Gatan) at 81,000x magnification, physical pixel size of 1.068 Å at a dose rate of 13.15 $e^{-}$ Å$^{-2}$ $s^{-1}$, exposure time of 4.23 s, corresponding to a total dose of 55.6 $e^{-}$ Å$^{-2}$ over 40 fractions.

Other datasets were collected in counted super-resolution mode on a Titan Krios G3 (FEI) operating at 300 kV with a BioQuantum imaging filter (Gatan) and K3 direct detection camera (Gatan) at 105,000× magnification, physical pixel size of 0.832 Å. Dose rates over 40 fractions for these data were as follows: *Cca*GldLM″, 22.2 $e^{-}$ Å$^{-2}$ $s^{-1}$, exposure time of 2.66 s, corresponding to a total dose of 59.1 $e^{-}$ Å$^{-2}$; *Sth*GldLM′ without fOM, 21.0 $e^{-}$ Å$^{-2}$ $s^{-1}$, exposure time of 2.97 s, corresponding to a total dose of 62.4 $e^{-}$ Å$^{-2}$; *Sth*GldLM′ with fOM, 20.6 $e^{-}$ Å$^{-2}$ $s^{-1}$, exposure time of 2.97 s, corresponding to a total dose of 61.2 $e^{-}$ Å$^{-2}$; *Swe*GldLM′, 21.4 $e^{-}$ Å$^{-2}$ $s^{-1}$, exposure time of 2.66 s, corresponding to a total dose of 56.9 $e^{-}$ Å$^{-2}$.

#### Cryo-EM data processing

##### *Cca*GldLM″

9,197,926 particles were extracted from 11,840 movies in 256 x 256-pixel boxes. After a round of reference-free 2d classification in SIMPLE, 4,607,771 particles were selected and reextracted in 412 x 412-pixel boxes. After another round of reference-free 2d classification in SIMPLE many classes had two particles visible due to the large box size and mask radius used. 2,009,669 particles with only one particle visible were selected and used to generate an *ab initio* model with C2 symmetry in SIMPLE. A larger selection of 4,282,288 particles, including doubleparticle classes, was made and rescaled and reextracted in 208 x 208-pixel boxes with a 1.648 Å pixel size.

The *ab initio* model was 60 Å low pass filtered and used as a reference for 3d classification with no symmetry applied with 3 classes and 7.5 ° sampling for 5 iterations. One class was selected, and extraneous density manually removed before use as a mask and reference (with 60 Å low pass filter) for another 3d classification of the whole dataset with 5 classes and 7.5 ° sampling for 15 iterations. One class with 1,236,725 particles was selected and the map used as reference (with 15 Å low pass filter) for a round of unmasked 3d classification with this class for 15 iterations with 4 classes and 7.5 Å sampling. One of the resulting classes had good density for the D2 domain but poor density for the transmembrane helices, another had good density for the transmembrane helices but very poor density for the D2 domain. A supervised 3d classification was used to sort all particles from the whole dataset into each of these two classes. 1,793,574 particles were sorted into the class with good D2 domain density (D2 class) and 2,488,741 into the class with good transmembrane helix density (TMH class). Once the transmembrane helices were clearly visible the map handedness was flipped to match the other structures.

A 3d classification using the D2 map from the previous classification as reference (with 15 Å low pass filter) was run on the D2 dataset with four classes and 7.5 ° sampling for 15 iterations then 3.75 ° sampling for 10 iterations. One class with 914,173 particles was selected and used for 3d autorefinement, producing a 3.7 Å map. The transmembrane helices were poorly defined, and the density could not be improved by micelle-focused refinements. A further 3D classification of these 914,173 particles with 4 classes and 7.5 ° sampling for 15 iterations yielded two classes totalling 595,559 particles with well-defined D2 domain density but poor TMH density and a class of 225,439 particles with improved TMH density but weakened D2 domain density. A periplasmic-domain focused refinement of the first class gave a 3.4 Å map with the micelle only visible at low contour level. We refer to this map and the atomic model built into it as *Cca*GldLM″$_{peri}$. Extensive processing of the second class in RELION 3.1 and cryoSPARC 2.15 did not produce a map in which both the D2 and TMH regions were well defined (7, 8).

Further 3d classification and refinement of the TMH class gave maps with distorted density due to a low proportion of side views. A new selection of 1,547,735 particles with a better distribution of views was made in SIMPLE and reextracted in 256 x 256-pixel boxes with a 0.832 pixel size. A 3d classification using the TMH class map with a 60 Å low pass filter as a reference was run with 4 classes for 15 iterations with 7.5 ° sampling. One class with 531,709 particles was selected and used for unmasked 3d autorefinement followed by masked 3d autorefinement. This yielded a 3.2 Å map but some distortions were still visible. The model of *Fjo*GldLM′ (PDB 6sy8) was used to generate a protein-only mask that was used for 3d classification without alignment. One class with 131,883 particles was selected and 3d autorefinement produced a 3.1 Å map without distortions. Bayesian particle polishing and further 3d classification without alignment focused on the periplasmic domains gave a final map at 3.0 Å from 77,223 particles. We refer to this map and the atomic model built into it as *Cca*GldLM″$_{TMH}$.

##### *Pgi*PorLM′

From the initial (K2 detector) dataset 403,648 particles from 5,268 movies were extracted in 256 x 256-pixel boxes. After two rounds of reference-free 2d classification in SIMPLE 328,271 particles were selected. A map of *Fjo*GldLM was low pass filtered to 60 Å and used as a reference for 3d classification in RELION 3.1 with 3 classes for 15 iterations with 7.5 ° sampling followed by 10 iterations with 3.75 ° sampling. One class with 151,667 particles was selected and used for 3d autorefinement, yielding a 6.4 Å map. This map was used as an initial reference (with 40 Å low pass filter) and mask for another round of 3d classification and 3d autorefinement, yielding a 5.0 Å map.

From the second (K3 detector) dataset 8,208,503 particles were extracted from 13,562 movies in 256 x 256-pixel boxes and subjected to two rounds of reference-free 2d classification in SIMPLE, from which 3,056,944 particles were selected. After another round of reference-free 2d classification in RELION 3.1 2,513,045 particles were selected. The map produced from the initial dataset was used to create a mask and initial reference (40 Å low pass filtered) for 3d classification with 4 classes and 7.5° sampling. One class with 633,283 particles was selected and used for 3d autorefinement without a mask. The resulting 4.9 Å map was used as mask and reference (with 15 Å low pass filter) for another round of 3d autorefinement, followed by Bayesian particle polishing and per-particle CTF refinement. This yielded a final map of 3.9 Å.

##### *Sth*GldLM′

From the initial (K2 detector) dataset 1,466,521 particles were extracted from 7,008 movies in 256 x 256-pixel boxes. After 2 rounds of reference-free 2d classification in SIMPLE 711,753 particles were selected. A map of *Fjo*GldLM was low pass filtered to 60 Å and used as a reference for 3d classification in RELION 3.1 with 4 classes for 15 iterations with 7.5 ° sampling then 10 iterations with 3.75° sampling. 1 class of 185,820 particles was selected and used for 3d autorefinement, yielding a 6.0 Å map. This map was used as reference (with 40 Å low pass filter) and mask for another round of 3d classification using the 711,753 particle selection with four classes for 15 iterations with 7.5 ° sampling then 10 iterations of 3.75 ° sampling. One class of 316,792 particles was selected and used for 3d autorefinement without and then with a mask, yielding a map of 5.4 Å. Attempts to improve this resolution were unsuccessful due to the poor view distribution of the dataset (Figure S6e).

From the sample with fluorinated octyl maltoside (K3 detector), 4,539,707 particles were extracted in 256 x 256-pixel boxes. After 2 rounds of reference-free 2d classification in SIMPLE 817,030 particles were selected, some of which seemed to show a string-like object in proximity to the detergent micelle. After another round of reference-free 2d classification in RELION 3.1 386,285 particles were selected. From the sample without fluorinated octyl maltoside (K3 detector), 9,203,748 particles were extracted from 22,420 movies in 256 x 256-pixel boxes. After two rounds of reference-free 2d classification in SIMPLE 3,135,006 were selected. After a further round of reference-free 2d classification in RELION 3.1 1,422,930 particles were selected. The two datasets were combined and used for 3d classification with 15 Å low pass filtered map from the previous dataset as an initial reference for 15 iterations with 7.5 ° sampling. The string-like object was not well resolved. One class with 749,660 particles was selected and used for 3d autorefinement, yielding a 3.7 Å map. Further refinement after Bayesian particle polishing and 3d classification focused on the transmembrane helices yielded a final 3.0 Å map from 394,678 particles.

*Swe*GldLM′

7,167,266 particles were extracted from 12,495 movies in 256 x 256-pixel boxes. After two rounds of reference-free 2d classification in SIMPLE 3,873,460 were selected. The *Sth*GldLMd1c map was used as a reference with a 60 Å low pass filter for 3d classification in

RELION 3.1 with 5 classes for 15 iterations with 7.5° sampling. One class with 1,513,439 classes was selected and used for 3d autorefinement, producing a 3.1 Å map with distortions due to lack of side views.

A harsher selection of 1,360,637 particles was made in SIMPLE with a greater proportion of side views retained. After 3d classification using the distorted map with 40 a Å low pass filter as a reference and four classes with 7.5° sampling for 15 iterations one class with 498,523 particles was selected. After 3d autorefinement a 3.5 Å map with some distortions visible in the periplasmic domains was produced. After periplasm-focused 3d classification without alignment a 3.5 Å map without periplasmic distortions was produced from 160,612 particles. Further 3d autorefinements after Bayesian particle polishing and a transmembrane helix-focused 3d classification produced a final 3.0 Å map from 111,727 particles.

### Supplemental Tables

**Table S1. Bacterial strains used in this study.**

| Strain | Genotype | Reference |
|---|---|---|
| ***E. coli*** | | |
| BL21 Star$^{TM}$ (DE3) | *$F^-$ ompT hsdS$_B$(r$_{B-}$, m$_{B-}$) gal dcm rne131 (DE3)* | Invitrogen |
| S17-1 | *Pro, res$^-$ hsdR17 (rK$^-$ mK$^+$) recA$^-$, RP4-2-Tc::Mu-Km::Tn7, Tp$^r$* | (4) |
| ***F. johnsoniae*** | | |
| UW101 | | (9) |
| Fl_082 | UW101 *ΔgldL* | (2) |
| Rhj_074 | UW101 *gldL*$_{64\text{-}74GSS}$ | This study |
| Rhj_076 | UW101 *gldL*$_{\Delta 64\text{-}74}$ | This study |

**Table S2. Plasmids used in this study**

| Plasmid | Description | Reference |
| --- | --- | --- |
| pT12_ SpaPQR3xFLAG | Encodes Salmonella enterica serovar Typhimurium SpaPQR operon with a C-terminal 3xFLAG tag on SpaR under the control of the E. coli rhaB promoter; ori cloDF13, $Kan^r$ | (1) |
| pMA-RQ_SthPKLMN | Synthetic plasmid with *S. thermophila* strain DSM 21410 genes RCX05217-05221 and 100 bp upstream and downstream in the pMA-RQ vector. Ori Col E1, $Amp^r$ | This study. Supplied by GeneArt (Invitrogen). |
| pRHJ012 | pGEM-T *gldL* | (2) |
| pRHJ113 | pMA-RQ_SthPKLMN with NcoI, NdeI and BamHI sites in GldL and GldM, respectively, removed by Quikchange mutagenesis | This study |
| pRHJ117 | pT12 *S. thermophila gldL gldM(1-229)-twinstrep* | This study |
| pRHJ118 | pT12 *P. gingivalis porL porM(1-227)-twinstrep* | This study |
| pRHJ170 | pT12 *S. wenxiniae gldL gldM(1-224)-twinstrep* | This study |
| pRHJ174 | pT12 *C. canimorsus gldL gldM(1-330)-twinstrep* | This study |
| pRHJ237 | pGEM-T $gldL_{64\text{-}74GSS}$ | This study |
| pRHJ238 | pGEM-T $gldL_{\Delta 64\text{-}74}$ | This study |
| pRHJ240 | Suicide plasmid used to introduce the E64-L74 to GSSGSSGSSGS codon change into *gldL*; $gldL_{64\text{-}74GSS}$ in pYT354 | This study |
| pRHJ241 | Suicide plasmid used to delete E64-L74 from *gldL*; $gldL_{\Delta 64\text{-}74}$ in pYT354 | This study |

**Table S3. Oligonucleotides used in this study.**

| Primer | Sequence 5'-3' | Used in the construction of |
|---|---|---|
| RHJ035 | AGGTATGCTTTCGGCGATGGGCGTTAGCAAG | pRHJ113, mutation site is underlined |
| RHJ036 | CTTGCTAACGCCCATCGCCGAAAGCATACCT | pRHJ113, mutation site is underlined |
| RHJ037 | GCATTCAGACTCTCCATGTGCTTTGCAGCAAGAGT | pRHJ113, mutation site is underlined |
| RHJ038 | ACTCTTGCTGCAAAGCACATGGAGAGTCTGAATGC | pRHJ113, mutation site is underlined |
| RHJ039 | ATTGTTTATCAATGAAGGATCAAGAGGTTGGCCATTGATGTAG | pRHJ113, mutation site is underlined |
| RHJ040 | CTACATCAATGGCCAACCTCTTGATCCTTCATTGATAAACAAT | pRHJ113, mutation site is underlined |
| RHJ162 | GAAAACCTGTACTTCCAGG | Used to linearise pT12 |
| RHJ163 | GGTGAATTCCTCCTGAATTTC | Used to linearise pT12 |
| RHJ168 | AAATTCAGGAGGAATTCACCATGGGTCATTATAGAAGATACAAGAAC | pRHJ118 |
| RHJ177 | AAATTCAGGAGGAATTCACCATGCCACTTATTGATGTAAACG | pRHJ117 |
| RHJ178 | TTATATGTTTACTTGCTAACGCCCAT | pRHJ117 |
| RHJ179 | GCGTTAGCAAGTAAACATATAAACATTAAATCAGAATAATTTAAAATGGCT | pRHJ117 |
| RHJ202 | CCTGGAAGTACAGGTTTTCACTGTTCACACGATAATCCC | pRHJ118 |
| RHJ203 | CCTGGAAGTACAGGTTTTCATCGGTAAACTTTAAATCGGTTTT | pRHJ117 |
| RHJ341 | CGGCCGCTCTAGAACTAGTGGATCCGCTGAGCCTGTATTTCGAG | pRHJ240 and pRHJ241 |
| RHJ342 | TACCGGGCCCCCCCTCGAGGTCGACCAGTACTTCACG | pRHJ240 and pRHJ241 |
| RHJ452 | AAATTCAGGAGGAATTCACCATGGCACAATCAAACAAAAC | pRHJ174 |
| RHJ471 | AAATTCAGGAGGAATTCACCATGGCAAAGAAAACAAAATTCAA | pRHJ170 |
| RHJ473 | CCCTGGAAGTACAGGTTTTCCTGGTCTAAGTTAACTACTGCC | pRHJ170 |
| RHJ481 | CCCTGGAAGTACAGGTTTTCAGGTTTAGGAATTGTTGAAAATACTTG | pRHJ174 |

| | | |
|---|---|---|
| RHJ617 | TCTGGGTCTTCAAGTGGCTCATCAGGAGCTAACGGACAAGCTAG | pRHJ237, mutation site is underlined |
| RHJ618 | GAGCCACTTGAAGACCCAGATGATCCATCCTCAACTGGTTCAAAAG | pRHJ237, mutation site is underlined |
| RHJ621 | AGTTGAGGATGCTAACGGACAAGCTAG | pRHJ238 |
| RHJ622 | GTCCGTTAGCATCCTCAACTGGTTCAAAAG | pRHJ238 |

### Supplemental Figures

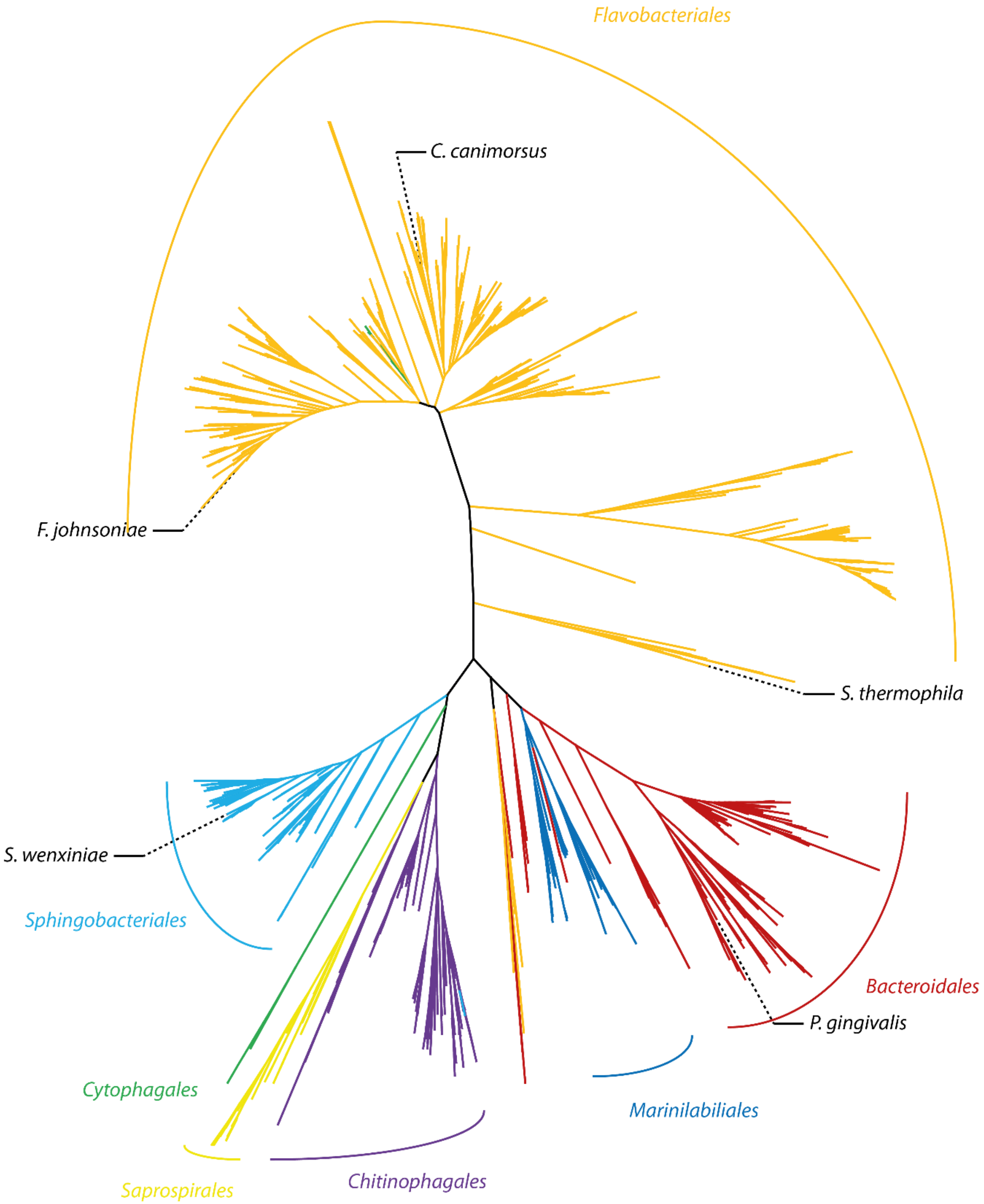

**Figure S1 Maximum likelihood phylogenetic tree of GldM sequences in the *Bacteroidetes* phylum.** Branches are colored by taxonomic order and the positions of proteins for which structures were determined are indicated.

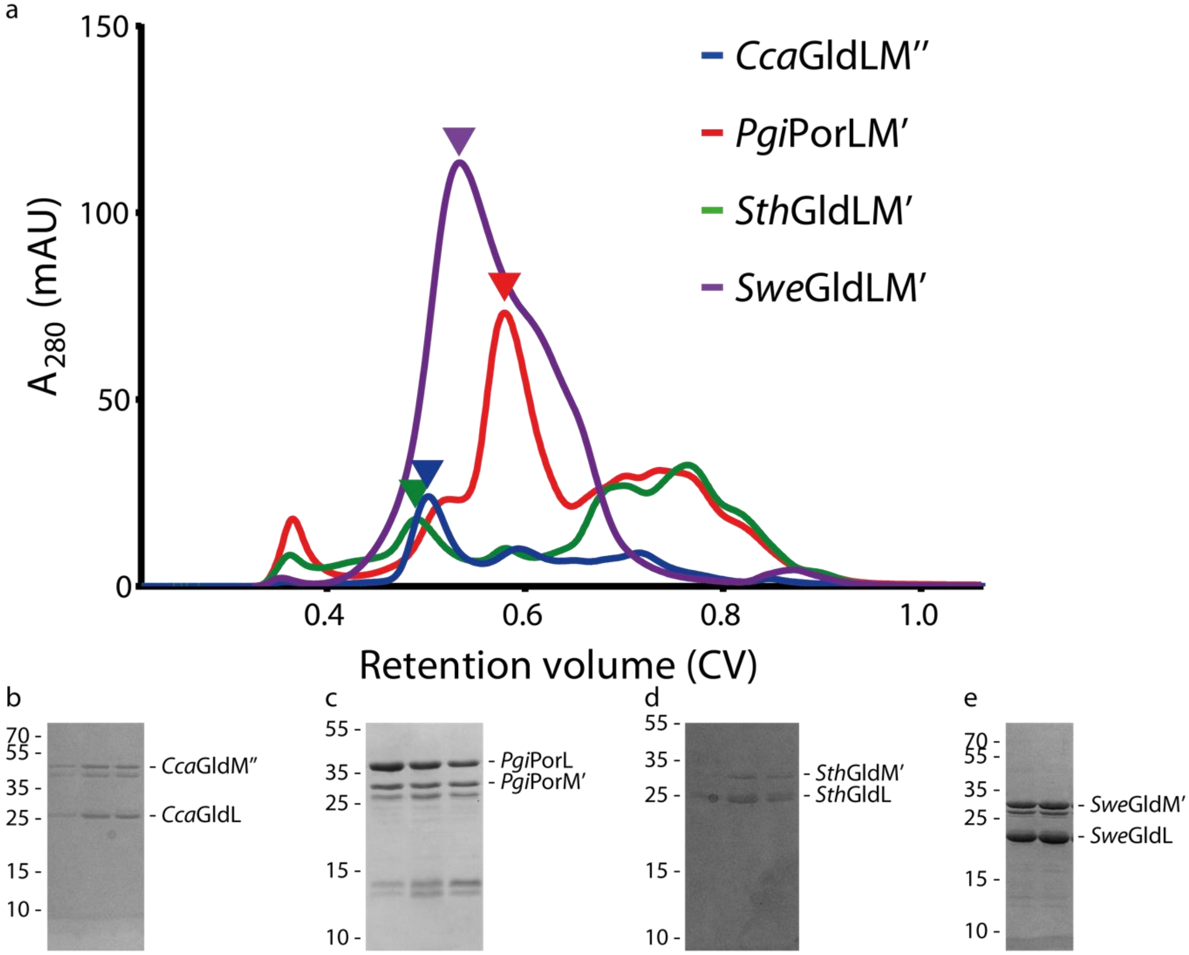


**Figure S2 Purification of GldLM'', PorLM', and GldLM' complexes. a** Size exclusion-chromatography traces for the indicated protein complexes. Proteins were analysed using a Superose 6 10/300 Increase column (Cytiva). The peaks used to prepare cryo-EM grids are indicated with arrowheads. **b-e** SDS-PAGE gels for the fractions used to make cryo-EM grids for the indicated protein complexes**.**

224

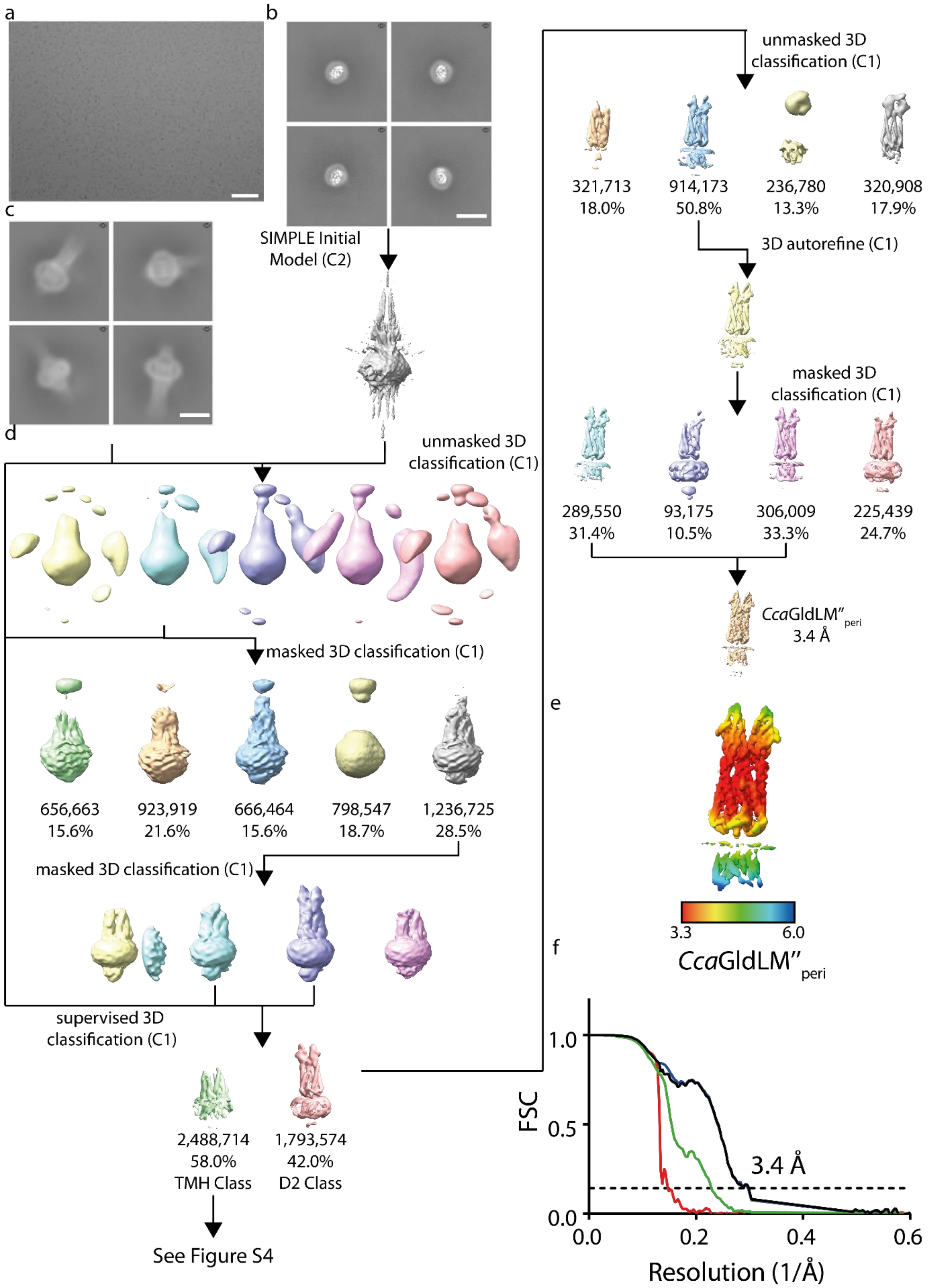
a
b
c
d
SIMPLE Initial Model (C2)
unmasked 3D classification (C1)
masked 3D classification (C1)
656,663 15.6%
923,919 21.6%
666,464 15.6%
798,547 18.7%
1,236,725 28.5%
masked 3D classification (C1)
supervised 3D classification (C1)
2,488,714 58.0% TMH Class
1,793,574 42.0% D2 Class
See Figure S4
unmasked 3D classification (C1)
321,713 18.0%
914,173 50.8%
236,780 13.3%
320,908 17.9%
3D autorefine (C1)
masked 3D classification (C1)
289,550 31.4%
93,175 10.5%
306,009 33.3%
225,439 24.7%
CcaGldLM''peri 3.4 Å
e
3.3
6.0
CcaGldLM''peri
f
FSC
1.0
0.5
0.0
3.4 Å
0.0
0.2
0.4
0.6
Resolution (1/Å)

**Figure S3 Data processing workflow for *Cca*GldLM″$_{\mathbf{peri}}$ map. a** Example micrograph for the *Cca*GldLM′′ sample collected at approximately 2.6 µm defocus. Scale bar 500 Å. **b** Representative 2D class averages used to produce the initial model. Scale bar 100 Å. **c** Representative 2D class averages used to produce the *Cca*GldLM″$_{peri}$ map. Scale bar 100 Å. **d** Data processing workflow for the *Cca*GldLM″$_{peri}$ map. The handedness of the maps was flipped between the left and right columns once helices were visible. **e** Local resolution estimates (in Å) for the sharpened *Cca*GldLM″$_{peri}$ map. **f** Fourier Shell Correlation (FSC) plot for the *Cca*GldLM″$_{peri}$ map. The resolution at the gold-standard cut-off (FSC = 0.143) is indicated by the dashed line. Curves: Red, phase-randomised; Green, unmasked; Blue, masked; black, MTF-corrected.

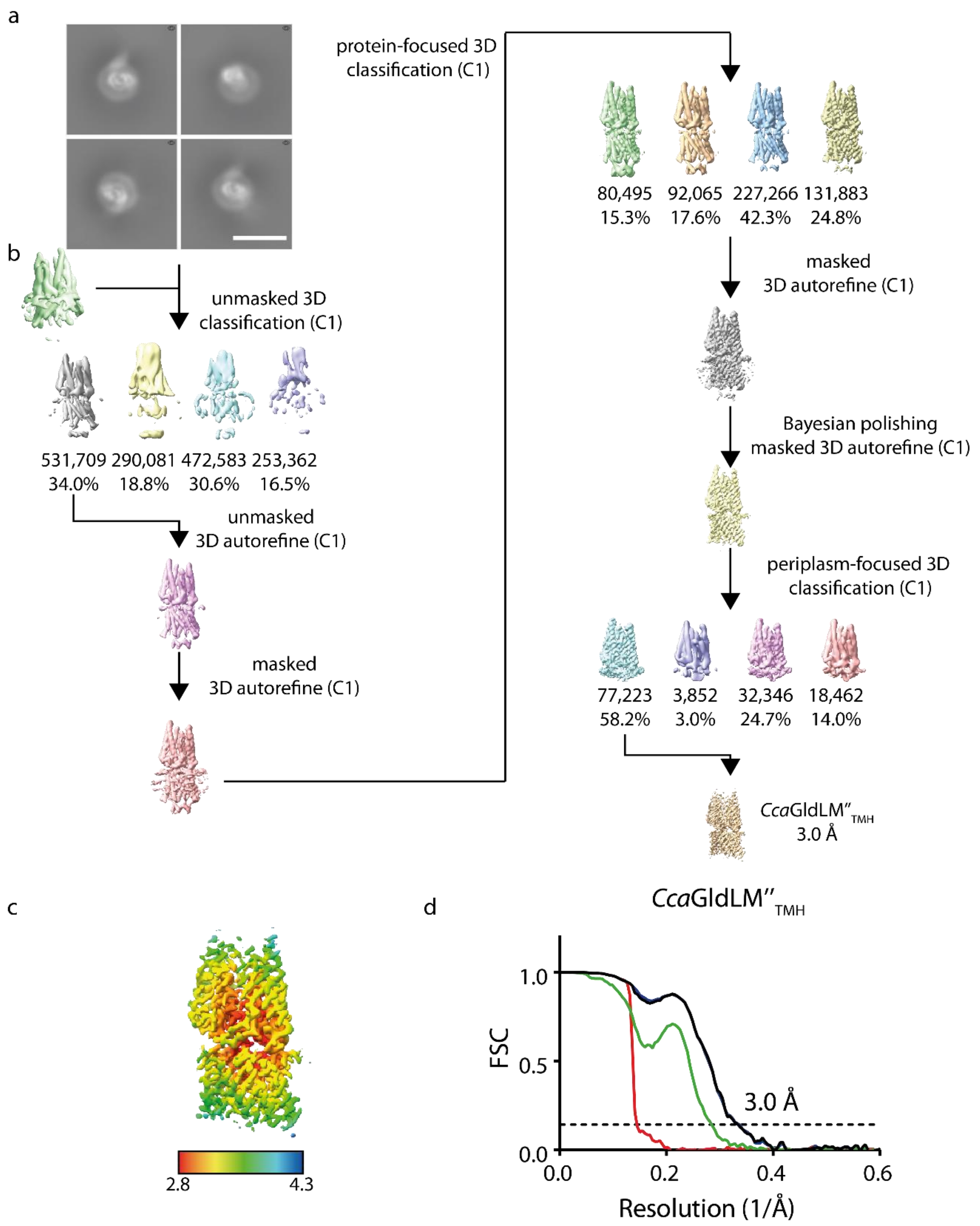


236 **Figure S4 Data processing workflow for *Cca*GldLM"$_{TMH}$ map. a** Representative 2D class averages used to produce the *Cca*GldLM"$_{TMH}$ map. Scale bar 100 Å. **b** Data processing workflow for the *Cca*GldLM"$_{TMH}$ map. **c** Local resolution estimates (in Å) for the sharpened *Cca*GldLM"$_{TMH}$

map. **d** FSC plot for the *Cca*GldLM″$_{TMH}$ map. The resolution at the gold-standard cut-off (FSC = 0.143) is indicated by the dashed line. Curves: Red, phase-randomised; Green, unmasked; Blue, masked; black, MTF-corrected.

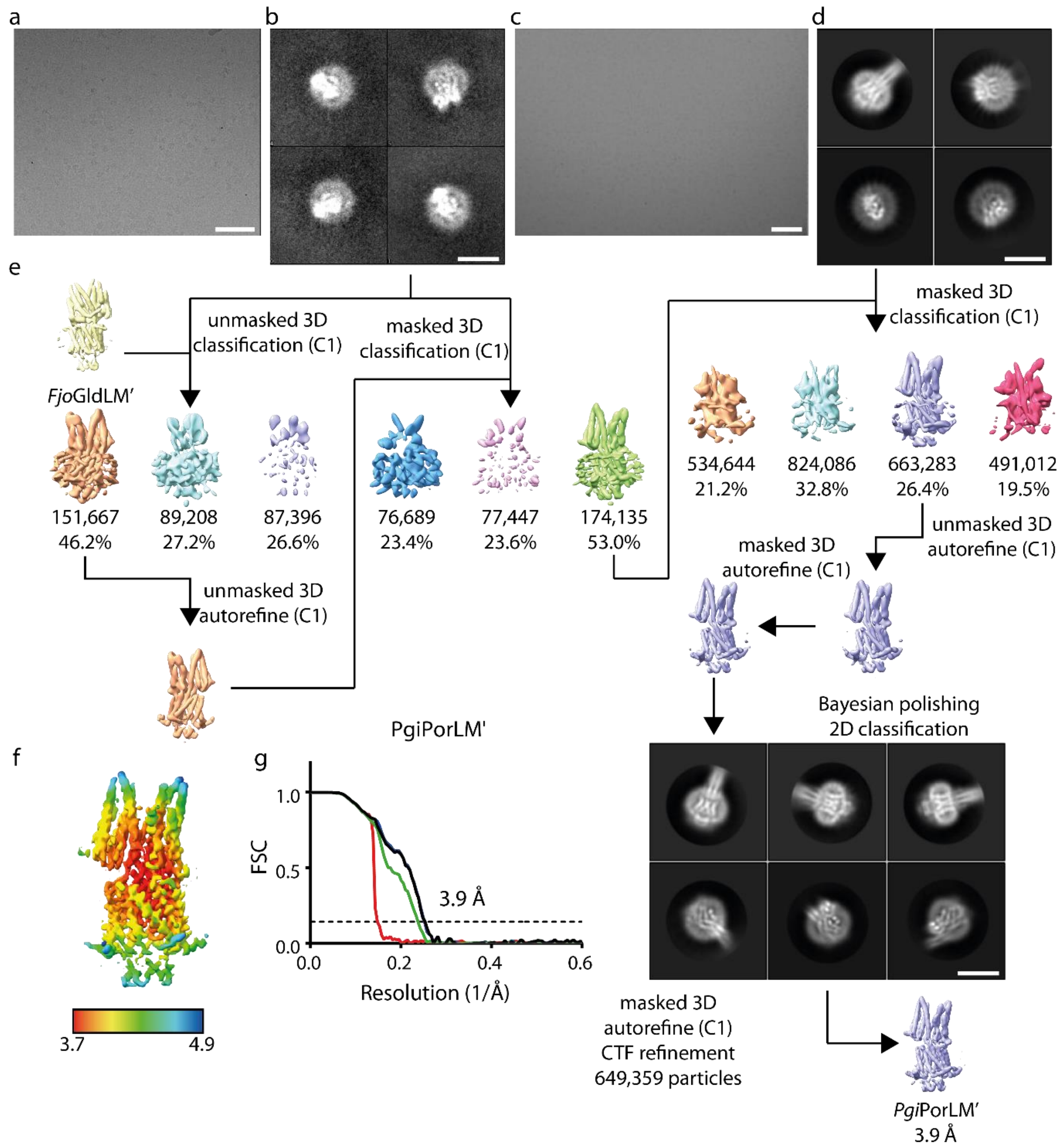


**Figure S5 Data processing workflow for *Pgi*PorLM' map. a** Example micrograph for the initial *Pgi*PorLM' dataset collected with a K2 detector at approximately 1.8 µm defocus. Scale bar 500 Å. **b** Representative 2D class averages from the K2 dataset used for initial processing. Scale bar 100 Å. **c** Example micrograph from the second *Pgi*PorLM' dataset collected with a K3 detector at approximately 1.8 µm defocus. Scale bar 500 Å. **d** Representative 2D class averages from the

K3 dataset used for final processing. Scale bar 100 Å. **e** Data processing workflow for the *Pgi*PorLM′ map. Scale bar on 2D class averages 100 Å. **f** Local resolution estimates (in Å) for the sharpened *Pgi*PorLM′ map. **g** Fourier Shell Correlation (FSC) plot for the *Pgi*PorLM′ map. The resolution at the gold-standard cut-off (FSC = 0.143) is indicated by the dashed line. Curves: Red, phase-randomised; Green, unmasked; Blue, masked; black, MTF-corrected.

254

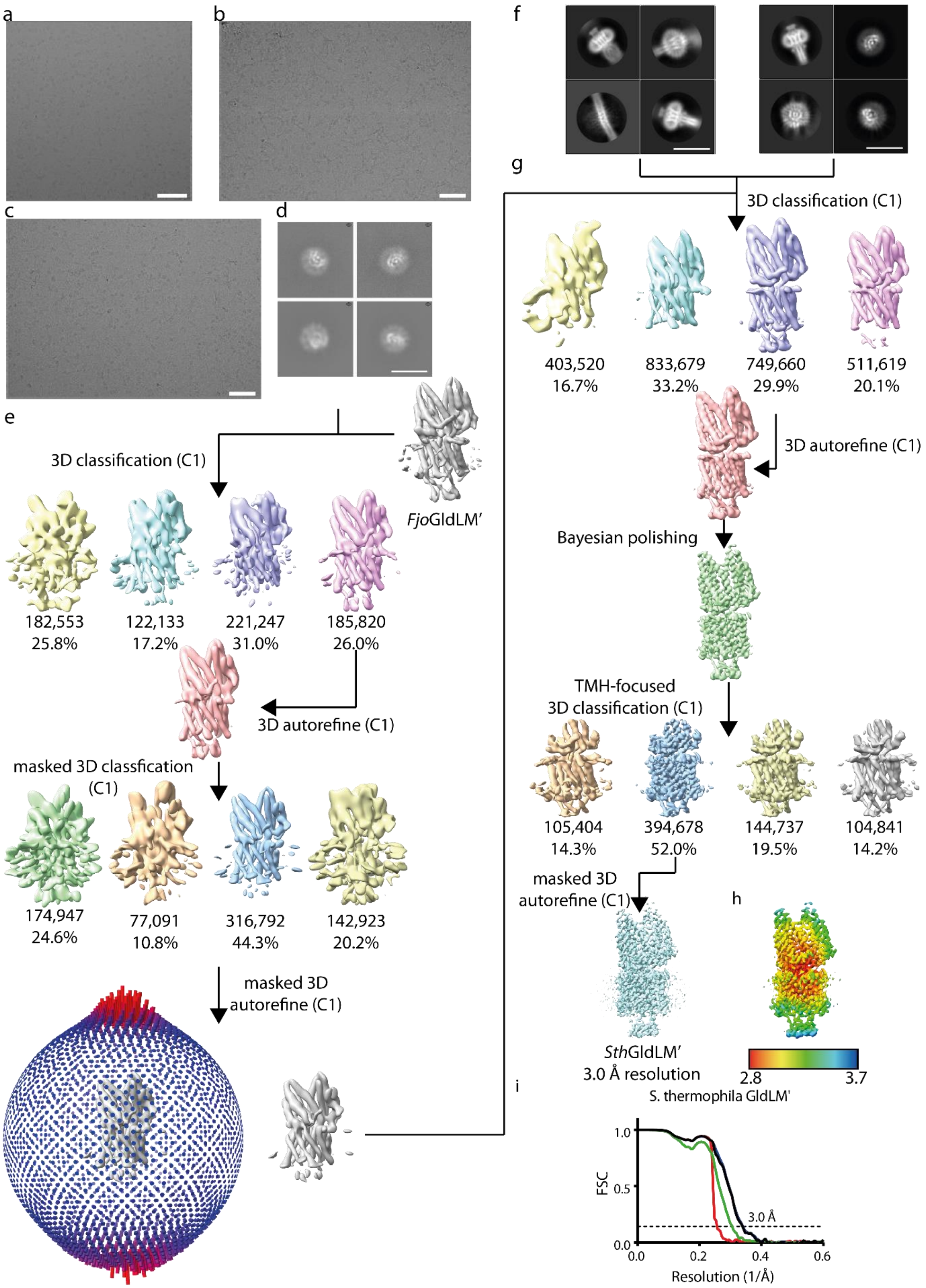
a
b
c
d
e
3D classification (C1)
FjoGldLM'
182,553
25.8%
122,133
17.2%
221,247
31.0%
185,820
26.0%
3D autorefine (C1)
masked 3D classfication (C1)
174,947
24.6%
77,091
10.8%
316,792
44.3%
142,923
20.2%
masked 3D autorefine (C1)
f
g
3D classification (C1)
403,520
16.7%
833,679
33.2%
749,660
29.9%
511,619
20.1%
3D autorefine (C1)
Bayesian polishing
TMH-focused 3D classification (C1)
105,404
14.3%
394,678
52.0%
144,737
19.5%
104,841
14.2%
masked 3D autorefine (C1)
SthGldLM'
3.0 Å resolution
h
2.8
3.7
i
S. thermophila GldLM'
FSC
1.0
0.5
0.0
3.0 Å
0.0
0.2
0.4
0.6
Resolution (1/Å)

**Figure S6 Data processing workflow for *Sth*GldLM′ map. a** Example micrograph for the initial *Sth*GldLM′ dataset collected with a K2 detector at a defocus of approximately 2.7 µm. **b** Example micrograph for the *Sth*GldLM′ + fOM dataset collected with a K3 detector at a defocus of approximately 1.7 µm. **c** Example micrograph for the *Sth*GldLM′ dataset collected with a K3 detector at defocus of approximately 1.7 µm. **d** Representative 2D class averages for the K2 dataset used in initial processing. **e** Data processing workflow for the initial dataset collected with a K2 detector. **f** Representative 2D class averages for the *Sth*GldLM′ + fOM (left) and *Sth*GldLM′ only (right) K3 datasets. **g** Data processing workflow for the combined datasets collected with a K3 detector. In the bottom-left panel each cylinder represents a view orientation, and the height of the cylinder corresponds to the number of views of that orientation. **h** Local resolution estimates (in Å) for the sharpened *Sth*GldLM′ map. **i** FSC plot for the *Sth*GldLM′ map. The resolution at the gold-standard cut-off (FSC = 0.143) is indicated by the dashed line. Curves: Red, phase-randomised; Green, unmasked; Blue, masked; black, MTF-corrected. Scale bars for micrographs 500 Å. Scale bars for 2D class averages 100 Å.

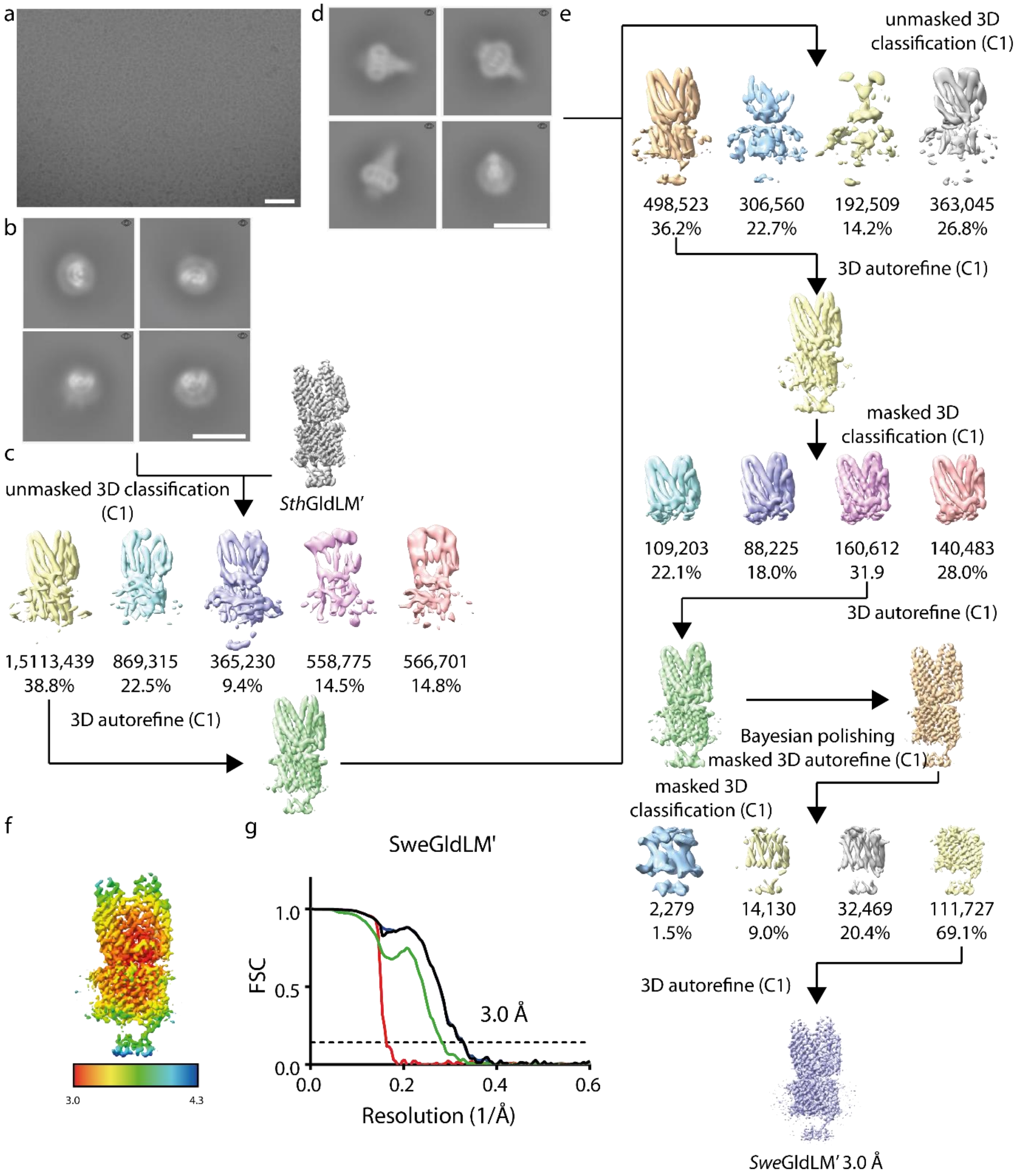


**Figure S7 Data processing workflow for *Swe*GldLM' map. a** Example micrograph for *Swe*GldLM' dataset collected with K3 detector at defocus of approximately 1.5 µm. Scale bar 500 Å. **b** Representative 2D class averages for initial particle selection. Scale bar 100 Å. **c** Data processing workflow for initial particle selection. **d** Representative 2D class averages for second, side-view focused particle selection. Scale bar 100 Å. **e** Data processing workflow for second particle selection. **f** Local resolution estimates (in Å) for the sharpened *Swe*GldLM' map. **g** Fourier Shell Correlation (FSC) plot for the *Swe*GldLM' map. The resolution at the gold-standard cut-off (FSC = 0.143) is indicated by the dashed line. Curves: Red, phase-randomised; Green, unmasked; Blue, masked; black, MTF-corrected.

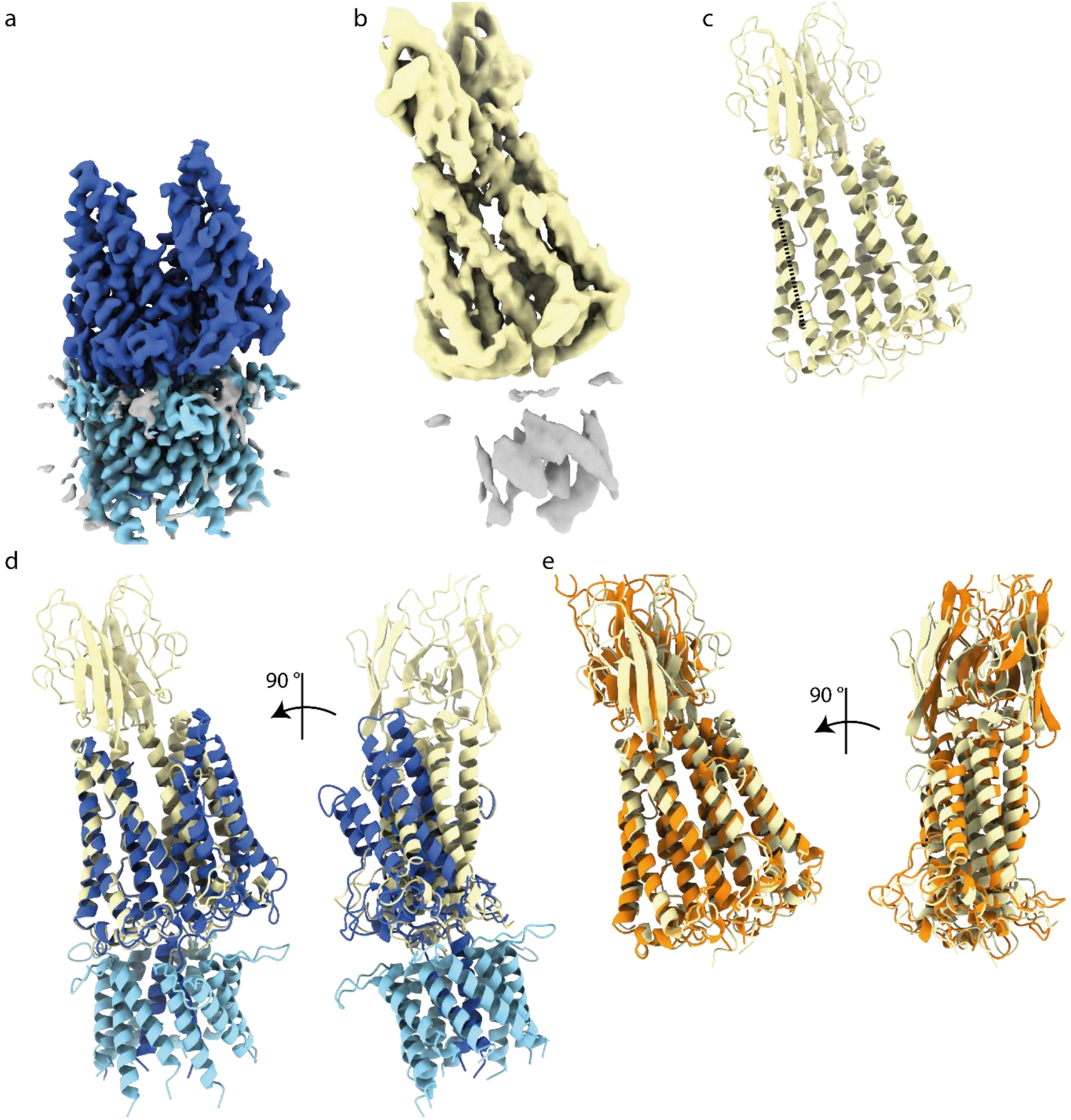


**Figure S8 Comparison of *Cca*GldLM″$_{\text{TMH}}$ and *Cca*GldLM″$_{\text{peri}}$ structures. a** *Cca*GldLM″$_{\text{TMH}}$ EM density map. GldL is colored light blue and GldM″$_{\text{TMH}}$ is colored dark blue. **b** *Cca*GldLM″$_{\textbf{peri}}$ EM density map. **c** Protein model of *Cca*GldLM″$_{\text{peri}}$. It was not possible to build protein into the transmembrane density. **d** Overlay of the *Cca*GldLM″$_{\text{TMH}}$ and *Cca*GldLM″$_{\text{peri}}$ structures showing the different relative orientations of the GldM D1 domains. Proteins are colored as in (a) and (c). **e** Overlay of *Cca*GldLM”$_{peri}$ (yellow) and *Fjo*GldM$_{peri}$ (PDB 6ey4)(orange) structures, showing the similarity in GldM D1 domain orientations. All maps and models in this figure were aligned to the second helix of *Cca*GldLM”$_{peri}$ Chain A, indicated with a dashed line in (c).

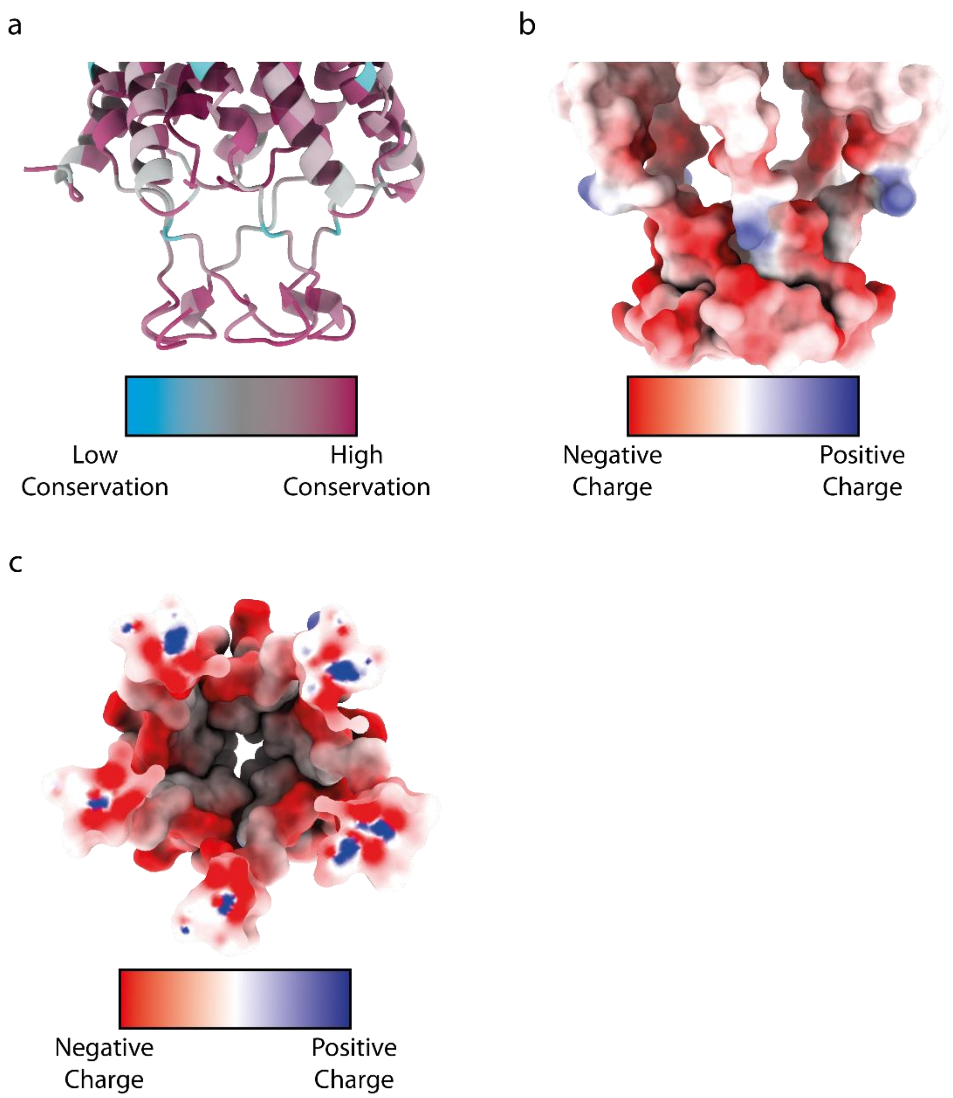


**Figure S9 Conservation and charge analysis for the cage region of *Sth*GldLM'. a** Sequence conservation analysis of the cage region of *S. thermophila* GldL using the program Consurf (10, 11). **b,c** Coulombic potential representation for the cage region of *S. thermophila* GldL. The view direction is parallel to the membrane (**b**) or from within the TMH bundle (**c**).For clarity, the first TMH and N-terminal residues are hidden.

#### References for the Supplemental Information


1. Dietsche T, Tesfazgi Mebrhatu M, Brunner MJ, Abrusci P, Yan J, Franz-Wachtel M, Schärfe C, Zilkenat S, Grin I, Galán JE, Kohlbacher O, Lea S, Macek B, Marlovits TC, Robinson CV, Wagner S. 2016. Structural and Functional Characterization of the Bacterial Type III Secretion Export Apparatus. PLOS Pathog 12:e1006071.

2. Hennell James R, Deme JC, Kjær A, Alcock F, Silale A, Lauber F, Johnson S, Berks BC, Lea SM. 2021. Structure and mechanism of the proton-driven motor that powers type 9 secretion and gliding motility. Nat Microbiol 6:221–223.

3. Zhu Y, Thomas F, Larocque R, Li N, Duffieux D, Cladière L, Souchaud F, Michel G, McBride MJ. 2017. Genetic analyses unravel the crucial role of a horizontally acquired alginate lyase for brown algal biomass degradation by *Zobellia galactanivorans*. Environ Microbiol 19:2164–2181.

4. Simon R, Priefer U, Pühler A. 1983. A broad host range mobilization system for in vivo genetic engineering: Transposon mutagenesis in Gram-negative bacteria. Bio/Technology 1:784–791.

5. McBride MJ, Kempf MJ. 1996. Development of techniques for the genetic manipulation of the gliding bacterium *Cytophaga johnsonae*. J Bacteriol 178:583–590.

6. Tartof KDA, Hobbs CA. 1987. Improved media for growing plasmid and cosmid clones. Bethesda Res Labs Focus 9:12.

7. Zivanov J, Nakane T, Forsberg BO, Kimanius D, Hagen WJH, Lindahl E, Scheres SHW. 2018. New tools for automated high-resolution cryo-EM structure determination in RELION-3. eLife 7:e42166.

8. Punjani A, Rubinstein JL, Fleet DJ, Brubaker MA. 2017. CryoSPARC: Algorithms for rapid unsupervised cryo-EM structure determination. Nat Methods 14:290–296.

9. McBride MJ, Braun TF. 2004. GldI is a lipoprotein that is required for *Flavobacterium johnsoniae* gliding motility and chitin utilization. J Bacteriol 186:2295–302.

10. Ashkenazy H, Abadi S, Martz E, Chay O, Mayrose I, Pupko T, Ben-Tal N. 2016. ConSurf 2016: an improved methodology to estimate and visualize evolutionary conservation in macromolecules. Nucleic Acids Res 44:W344–W350.

11. Landau M, Mayrose I, Rosenberg Y, Glaser F, Martz E, Pupko T, Ben-Tal N. 2005. ConSurf 2005: The projection of evolutionary conservation scores of residues on protein structures. Nucleic Acids Res 33:W299–W302.
